# 8 Å structure of the cytoplasmic ring of the *Xenopus laevis* nuclear pore complex solved by cryo-EM and AI

**DOI:** 10.1101/2021.11.10.468011

**Authors:** Linhua Tai, Yun Zhu, He Ren, Xiaojun Huang, Chuanmao Zhang, Fei Sun

**Affiliations:** National Key Laboratory of Biomacromolecules, CAS Center for Excellence in Biomacromolecules, Institute of Biophysics, Chinese Academy of Sciences, Beijing 100101, China; The Ministry of Education Key Laboratory of Cell Proliferation and Differentiation and the State Key Laboratory of Membrane Biology, College of Life Sciences, Peking University, Beijing 100871, China; Center for Biological Imaging, Institute of Biophysics, Chinese Academy of Sciences, Beijing 100101, China; University of Chinese Academy of Sciences, Beijing 100049, China; Bioland Laboratory (Guangzhou Regenerative Medicine and Health Guangdong Laboratory), Guangzhou, Guangdong 510005, China

## Abstract

As one of the largest protein complexes in eukaryotes, the nuclear pore complex (NPC) forms a conduit regulating nucleocytoplasmic transport. Here, we determined 8 Å resolution cryo-electron microscopic (cryo-EM) structure of the cytoplasmic ring (CR) from the *Xenopus laevis* NPC. With the aid of AlphaFold2, we managed to build a most comprehensive and accurate pseudoatomic model of the CR to date, including the Y complexes and flanking components of Nup358, Nup214 complexes, Nup205 and Nup93. Comparing with previously reported CR model, the Y complex structure in our model exhibits much tighter interactions in the hub region mediated by α-solenoid domain in Nup160 C-terminus. Five copies of Nup358 are identified in each CR subunit to provide rich interactions with other Nups in stem regions of Y complexes. Two copies of Nup214 complexes lay in a parallel pattern and attach to the short arm region of Y complexes towards the central channel of NPC. Besides, the structural details of two copies of Nup205 on the side of the short arm region and one copy of Nup93 on the stem region of Y complexes in each CR subunit are also revealed. These in-depth novel structural features represent a great advance in understanding the assembly of NPCs.

## Introduction

In eukaryotes, the double layer inner nucleus membrane (INM) and outer nucleus membrane (ONM) form the nuclear envelope (NE) to enclose nucleus to store genetic materials ^1,2^. To make the bidirectional transport possible between cytoplasm and nucleoplasm, nuclear pore complex (NPC) forms a conduit regulating nucleocytoplasmic transport. NPC is one of the biggest protein complexes throughout eukaryotes ^1,3,4^, and its building blocks are named as Nucleoporins (Nups). In fungi, NPC consists of ~500 Nups with molecular weight of ~66 MDa, while in higher eukaryotes, NPC consists of ~1000 Nups with molecular weight of ~120 MDa. These Nups arrange into a roughly eight-fold symmetrical assembly around a central channel perpendicular to the NE where the transportation occurs ^5–9^. NPC can be divided into four scaffolding rings and several attachments, including cytoplasmic ring (CR), inner ring (IR), nuclear ring (NR), luminal ring (LR), cytoplasmic filament, nuclear basket, and permeability barrier formed by phenylalanine-glycine (FG) rich Nups ^10–13^

Detailed structural information is necessary for mechanistic understanding of NPC functions, but it has long been hindered by the enormous size and high dynamicity of NPC. Cryo-electron tomography (cryo-ET) along with subtomogram averaging (STA) has been applied to reach ~2 nm resolution of NPC structures from multiple species, including *Homo sapiens* (*H. sapiens*), *Xenopus laevis* (*X. laevis*), *Chlamydomonas reinharadtii* (*C. reinharadtii*), *Schizosaccharomyces pombe* (*S. pombe*), and *Saccharomyces cerevisiae* (*S. cerevisiae*) ^13–19^. Based on these studies, the basic architectures of CR, IR and NR have been solved by rigidly docking of crystal structures of several Nups ^1,20^, such as the model reported by Lin et al. in 2016 (aliased as 2016-model) ^20^. In these scaffold rings, eight asymmetric units (or named as subunits) lay in a head to tail fashion to form the backbones. In outer rings (NR and CR), the backbones in each subunit are formed by one or two Y-shaped complexes, also known as Nup84 complex in fungi or Nup107-160 complex in vertebrates ^13,21–24^. In Y complex of vertebrates, Nup85, Nup43 and Seh1 form the short arm region, Nup160 and Nup37 form the long arm region, Sec13, Nup96, Nup107 and Nup133 form the stem region ^14^.

As a major member of NPC scaffold rings, the CR is essential for building and maintaining NPC structures. It provides docking sites for cytoplasmic filaments to regulate importin α/β dependent nucleocytoplasmic transport and messenger ribonucleoprotein (mRNP) exporting ^25,26^. Thus, in addition to the Y complex scaffold, CR has several unique components like Nup358 complex and Nup214 complex. However, the structural characteristics of these important CR components remain elusive, including the exact copy numbers, locations, and interactions. In 2020, cryo-electron microscopic (cryo-EM) single particle analysis (SPA) was applied to reveal the detailed structure of *X. laevis* NPC CR, reaching a highest resolution of 5.5 Å for most rigid part of CR (aliased as 2020-model) ^27^. However, due to the preferred orientations in sample preparation and lack of an accurate starting model, it’s still very hard to obtain a reliable pseudoatomic model of the CR.

Most recently, we developed an improved cryo-EM SPA method to solve the problem of preferred orientations, and determined the significantly improved structure of the NR of the *X. laevis* NPC with an isotropic resolution around 8 Å ^28^. Meanwhile, with the aid of the highly accurate protein structure prediction tool AlphaFold2 ^29^, we built the most complete pseudoatomic model of the NR and revealed multiple previously uncharacterized structural features. Here, using similar approaches, we determined the CR structure of the *X. laevis* NPC with an isotropic subnanometer resolution, and built the most complete and accurate pseudoatomic model of the CR to date. According to this significantly improved model, we identified multiple structural features in CR subunit, including tight interactions in the Y complex hub mediated by Nup160, five copies of Nup358 warped around the stem region, two copies of Nup214 complexes attached to the short arm region, two copies of Nup205 on the side of the arm region, single copy of Nup93 bridging the stem region. Our results improve the understanding of detailed assembly and functions of NPC.

## Results

### Structure determination of CR subunit of *X. laevis* NPC

Since NPCs are naturally perpendicular to the NE, imaging NPCs directly on native NE will bring significant problem of preferred orientations in the map reconstructions. As reported previously ^28^, we combined the strategy of imaging NPCs on tilted NE (tilt view) and on the edge of folded back NE (side view) to solve this problem (Extended Data Fig. 1). Using this approach, the reconstruction in core region of CR subunit reaches 8 Å isotropic resolution, with highest local resolution of 7.2 Å (Extended Data Fig. 2). Meanwhile, the whole CR subunit and CR Nup358 region reached the isotropic resolution of 8.7 Å and 8.9 Å, respectively (Extended Data Fig. 2). By investigate the directional resolution using three-dimensional Fourier shell correlation (3D-FSC) estimates ^30^, the sphericity values of these reconstructions range from 0.91 to 0.95, suggesting no significant anisotropy in the cryo-EM maps (Extended Data Fig. 2).

Then we utilized AlphaFold2 to predict the full-length model for each Nup in CR as the starting model for refinement (Extended Data Fig. 3, 4) ^31^. Based on the improved density map, sequential molecular dynamics flexible fitting (MDFF) and manual refinement were both used to build a total of 15 different Nups, 32 components into each CR subunit (Fig. 1A-B, Extended Data Fig. 3). We noticed that the refined model of each CR Nup according to the local density is basically similar with starting model (Extended Data Fig. 4), proving the high accuracy of the initial models generated by AlphaFold2. As the most comprehensive and accurate pseudoatomic model of the CR to date, there are 22372 residues in each CR subunit in our model, which extends 102% comparing with 2016-model (11064 residues) and 52.4% comparing with 2020-model (14683 residues) ^20,27^. Since the structures of β-propeller domains of CR Nups have been well studied in previous studies ^1,8,22,32,33^, these extensions are mainly located on α-solenoid domains of Y complex Nups and Nup93, Nup205, Nup358 and Nup214 complex, which will be discussed below (Fig. 1B-C).

**Figure 1.**
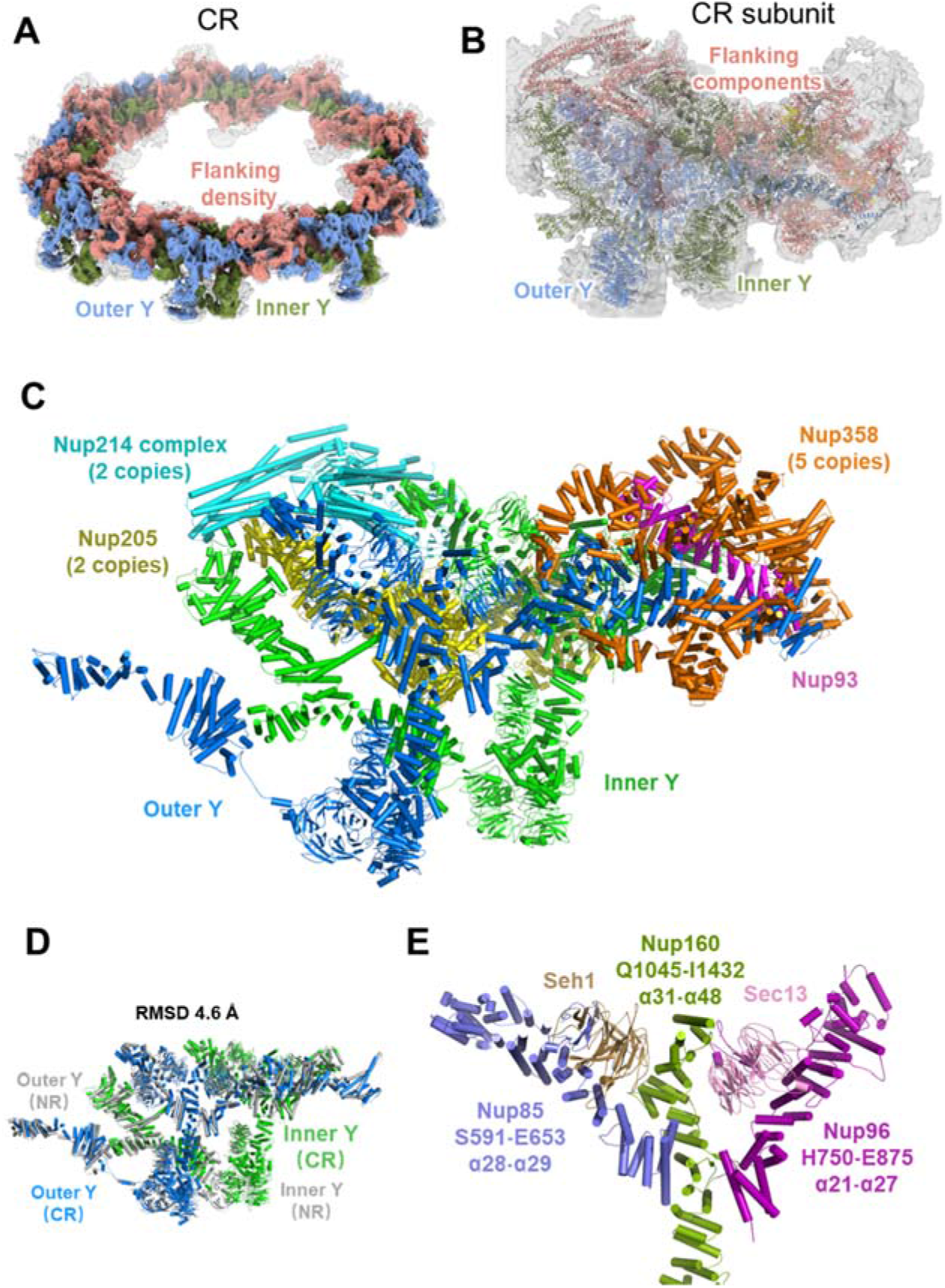
The more complete pseudoatomic model of the CR from the *X. laevis* oocyte NPC. (**A**) Overall view of the *X. laevis* NPC CR structure, displaying the inner and outer Y complexes in each asymmetric unit, as well as the densities other than the Y complexes. The inner Y complexes are colored in olive, the outer Y complexes are in cornflower blue, and the extra densities are in coral. (**B**) Model-map overlay of the NPC CR subunit. The map density is displayed in grey with transparency. Models of the inner Y complex, outer Y complex and extra densities are in the same colors as in (**A**). (**C**) Model display of Nups corresponding to two Y complexes, two Nup214 complexes, two Nup205 and five Nup358 in the CR subunit, coloured differently. (**D**) RMSD between two Y complexes within CR or NR subunit. (**E**) Y complex hub region in CR asymmetric unit, showing interactions of C terminals of Nup85, Nup160, Nup96 and Seh1, Sec13.

### Enhanced interactions of Y complexes in the hub region

Most recently, we reported the NR structure of *X. laevis* NPC, in which a total of 21 components and 15622 residues were built in each NR subunit ^28^. After comparing the more accurate models of NR and CR subunits, we found that besides the flanking components unique for each ring, like ELYS for NR and Nup358 or Nup214 complex for CR, their Y complex scaffold shares very similar architecture. The root-mean-square error (RMSD) value for double Y complexes in NR and CR is 4.6 Å (Fig. 1D), and no large shifts are found for all the domains in Y complex Nups. For individual Y complex, the RMSD values for inner Y and outer Y in CR, the inner Y in NR and CR, the outer Y in NR and CR are 5.6 Å, 4.8 Å and 3.8 Å, respectively (Extended Data Fig. 5). The only significant differences are found in comparison of CR inner and outer Y complexes, while the inner Nup133 has a shift of ~8 nm at the C-terminal domain (CTD). This shift should be related to the shorter circumference for eight inner Y complexes than outer Y complexes when forming a concentric ring of CR. The consistency of the scaffold structure in NR and CR agrees well with previous report ^13,14,20,34^.

In previous report, we have identified several novel interactions according to the most complete model of Y complex in NR ^28^. In this study, we also identified these interaction features in Y complex of CR. Briefly, in the Y complex hub region, the CTD of Nup160 (Q1045-I1432) recruits Seh1, Sec13, Nup85 and Nup96 to form an interaction network for stabilizing Y complex (Fig. 1E). It suggested that Nup160 plays a central role in the assembly and stabilizing Y complexes in both NR and CR. Moreover, it is worth noting that the local density map for N-terminal domain (NTD) of Nup160 in CR exhibits lower local resolution comparing to that in NR (Extended Data Fig. 2) ^28^, indicating the larger dynamic for the long arm region of Y complex in CR. This kind of dynamic of CR may be due to two reasons. On the one hand, in certain conditions like energy depletion, constriction may happen on the CR region of NPC ^19^. On the other hand, CR has only 32 β-propeller domains (16 from Nup160 and 16 from Nup133) anchored onto the NE, while NR have 8 or 16 more (from ELYS) to enhance the stability of Y complexes onto the membrane and increase the local stability of Nup160 NTD ^14^.

### Five copies of Nup358 reside in each CR subunit

Nup358, as the largest Nup in vertebrates, plays an essential role in biological functions of NPC through its multiple domains. It was known that Nup358 contains an α helical region in the NTD (Fig. 2A), followed by multiple domains separated by unstructured regions, including Ran-binding domain, Zinc finger domain, E3 ligase domain and Cyclophilin domain ^1,27,35–37^. Recent studies identified that the density corresponding to Nup358 looks like several clamps near the stems of Y complexes ^14,35^, and the copy number of Nup358 in each CR subunit may be 2 or 4 ^27,37^. By using AlphaFold2, we predicted the structure of Nup358 NTD, then found that this clamp-shaped structure could fit well in the local density of previously assigned location for Nup358 (Extended Data Fig. 3I). Strikingly, a total of 5 copies of Nup358 NTD could be well modeled into this region, suggesting that there should be at least 5 Nup358 proteins stably bound to the stem region of Y complexes in each CR subunit (Fig. 2B-C). The identification of 5 copies of Nup358 in each CR subunit could be also confirmed by fitting our CR model into the reported NPC structure from Hela cell, in which the density for all the five Nup358 proteins is obvious (Extended Data Fig. 6) ^14^.

**Figure 2.**
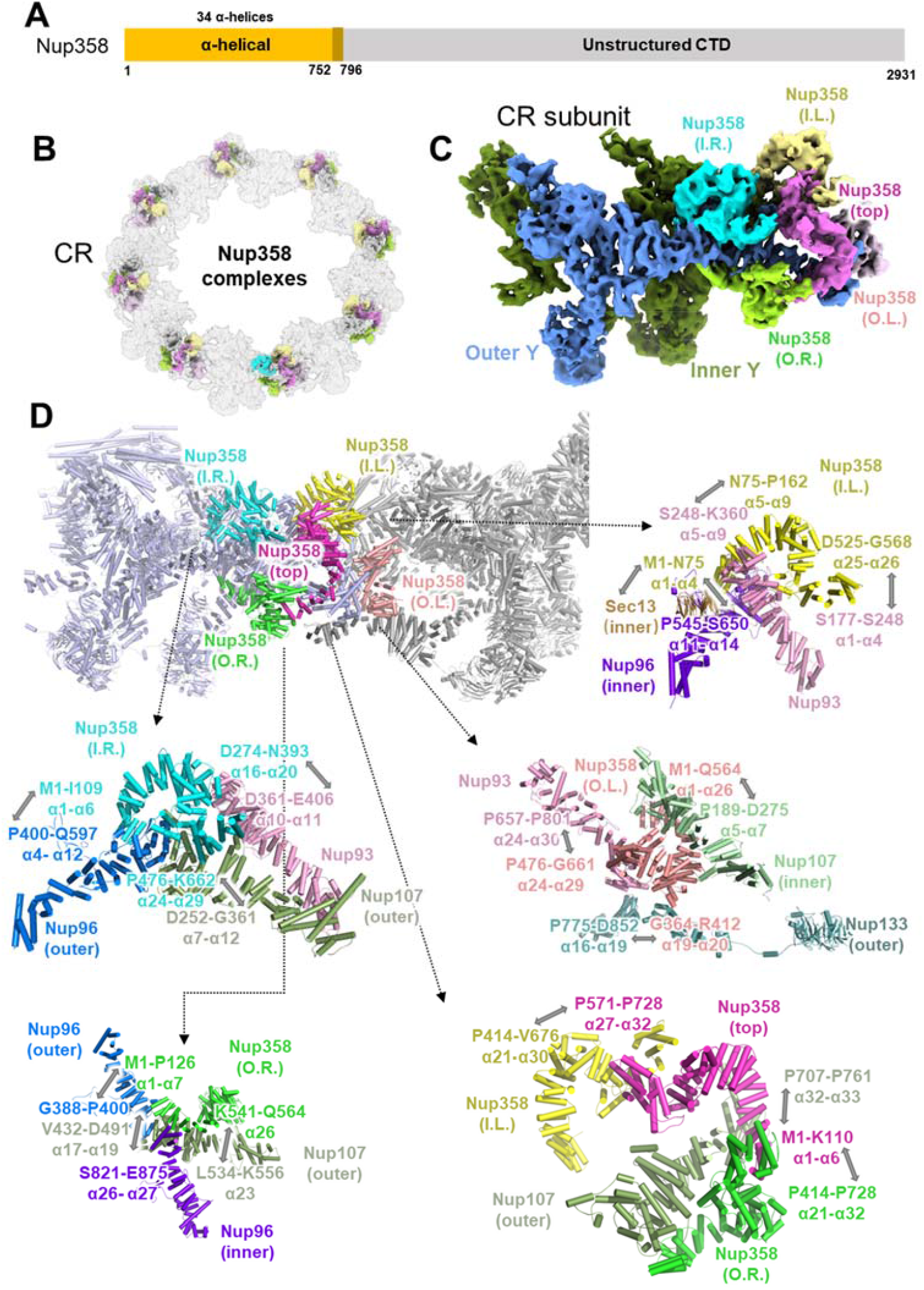
The structures and interaction details of 5 copies of Nup358 in each CR subunit. (**A**) Domain assignment of modeled part of Nup358. (**B**) Location of 40 copies Nup358 in CR, while the 5 copies of Nup358 in each subunit were colored differently. (**C**) Location of 5 copies of Nup358 in CR subunit, colored differently. (**D**) Interactions of 5 copies of Nup358 with surrounding Nups.

To distinguish each Nup358, we named them as inner-left, inner-right, outer-left, outer-right and top one according to their spatial locations, assuming the observer stands inside the nuclear channel (Fig. 2C). The two outer Nup358 proteins located farther from the nuclear channel than the inner ones, while the top Nup358 situated on top of the other four copies. These Nup358 proteins have rich contacts with surrounding CR Nups in different ways. Take inner-right Nup358 as an example, its α1 to α6 helices (M1 to I109) interact with α4 to α12 helices (P400 to Q597) of outer Nup96, α16 to α20 helices (D274 to N393) interact with α10 to α11 helices (D361 to E406) of Nup93, α24 to α29 helices (P476 to K662) interacts with α7 to α12 helices (D252 to G361) of outer Nup107 (Fig. 2D). The other four Nup358 have similar rich interactions (Fig. 2D). Briefly, outer-right Nup358 bind to outer Nup107 and inner/outer Nup96; inner-left Nup358 bind to Sec13, inner Nup96 and Nup93; outer-left Nup358 bind to inner Nup107, outer Nup133 and Nup93; top Nup358 binds to outer Nup107, inner-left Nup358 and outer-right Nup358. On the whole, the top Nup358 seems to act as a lid to cover the rest Nup358 proteins, and the later ones form direct and extensive interactions with inner and outer Y complex to stabilize the stem region. Moreover, the anchoring of these Nup358 NTDs onto CR will facilitate the other domains of Nup358 or other Nup358 related proteins to perform the proper biological functions at the right locations.

### Two Nup214 complexes lay in parallel in each CR subunit

As another major component of cytoplasmic filament, Nup214 complex is believed to form a mRNP export platform on the cytoplasmic face of NPC and coordinate the mRNP remodeling process to ensure the unidirectional transportation process ^38,39^. However, the structural details of Nup214 complex remains elusive to date, including the exact copy number, its relative locations to Y complex and the pseudoatomic model ^27^. The Nup214 complex is regarded to have at least three major components: Nup214, Nup88 and Nup62 in vertebrates (Fig. 3A), or Nup159, Nup82 and Nsp1 in fungi. It was reported that Nup159 complex may form a P-shaped homodimer configuration ^17,38,39^, but this kind of structure was not found in the CR structure of *X. laevis* ^27^.

**Figure 3.**
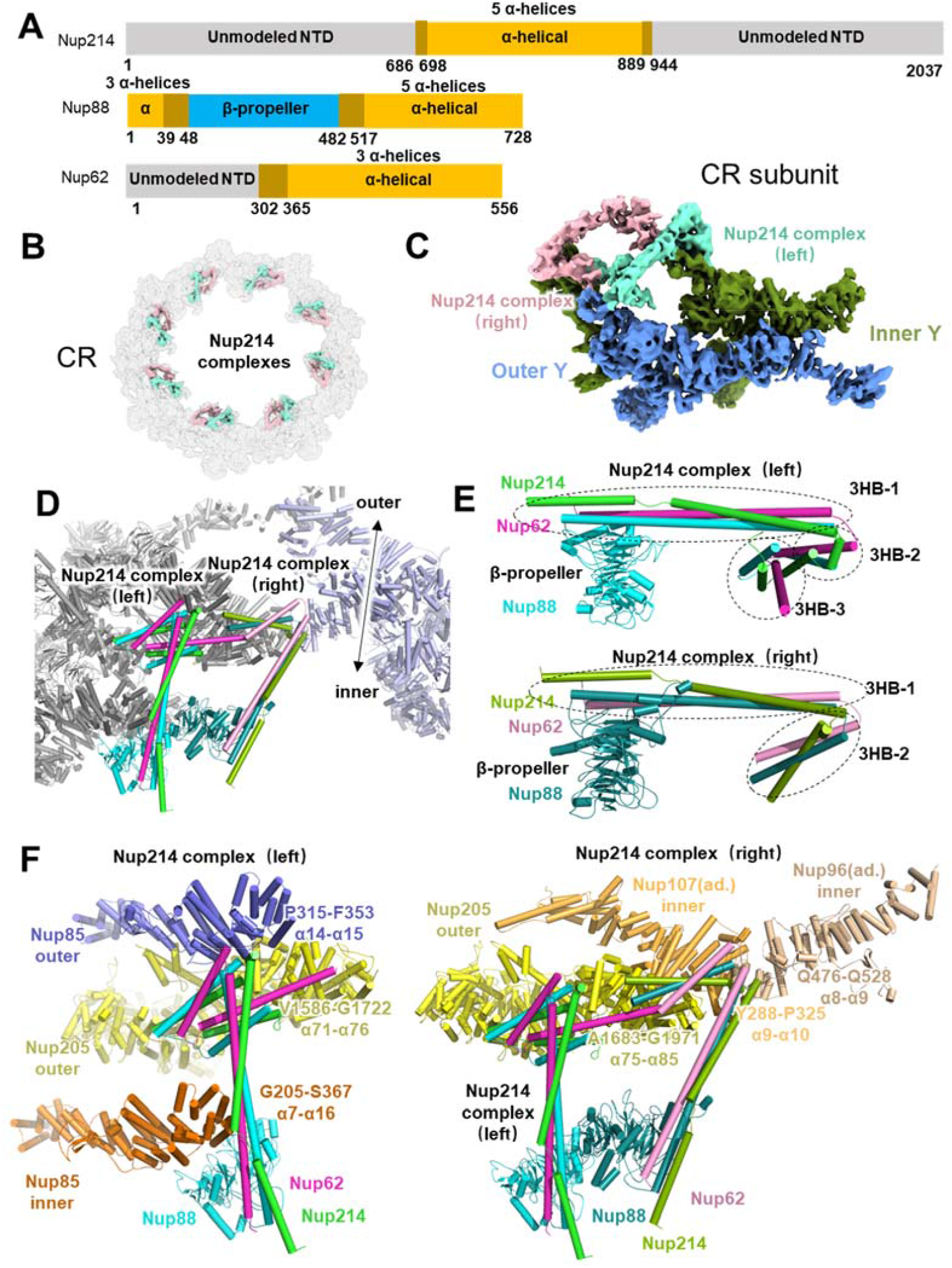
The structures and interaction details of 2 copies of Nup214 complex in each CR subunit. (**A**) Domain assignment of modeled part of Nup214, Nup88 and Nup62. (**B**) Location of 16 copies Nup214 complex in CR, Nup214 complex in each subunit were colored differently. (**C**) Location of 2 copies of Nup214 complex in CR subunit. The left Nup214 complex was colored in chartreuse, right Nup214 complex was in light pink. (**D**) Model of left and right Nup214 complex attached to two CR subunits. (**E**) Domain display of left and right Nup214 complexes. (**F**) Interaction sites of left and right Nup214 complexes.

Using AlphaFold2, we predicted the complex structure of Nup214, Nup88 and Nup62, and found that these three Nups could form a rake-shaped conformation (Extended Data Fig. 3J), including a β-propeller domain as the handle, a helix bundle as the body and two helix bundles as the head. Then we fitted this rake-shaped structure into the flanking density above the short arms of two Y complexes and found that there should be two copies of Nup214 complexes in each CR subunit (Fig. 3B-C). To distinguish these two complexes, we named them as left one and right one according to their spatial locations, assuming the observer stands inside the nuclear channel (Fig. 3D). In the left Nup214 complex, the rake handle is made up of β-propeller domain of Nup88, while the unusually long rake body is made up of three helices bundle (3HB) from Nup214, Nup88 and Nup62, named as 3HB-1. For the rake head, there are two 3HB structures, 3HB-2 and 3HB-3, while every helix in the 3HBs comes from different Nups. The right Nup214 complex has basically the same conformation as the left one, except for the missing 3HB-3 domain (Fig. 3E).

With this much improved model of those two Nup214 complexes in CR subunit, we found several important interaction features. For left Nup214 complex, the β-propeller of Nup88 and 3HB-1 domain binds to the α7 to α16 helices (G205 to S367) of inner Nup85. Then 3HB-2 and 3HB-3 make close contact with α14 to α15 helices (P315 to F353) of outer Nup85, and also α71 to α76 helices (V1586 to G1722) of outer Nup205 (Fig. 3F). For right Nup214 complex, the β-propeller of Nup88 binds to the left Nup214 complex, and its 3HB-1 and 3HB-2 domains connects with α75 to α85 helices (A1683 to G1971) of outer Nup205, α9 to α10 helices (Y288 to P325) of inner Nup107 in adjacent CR subunit, α8 to α9 helices (Q476 to Q528) of inner Nup96 in adjacent CR subunit (Fig. 3F). It seems that the two Nup214 complexes help to stabilize the two short arm regions of Y complexes in CR subunit and contribute to the head-to-tail fashion of adjacent CR subunits. Moreover, the NTDs of both left and right Nup214 complexes point to the nuclear channel, allowing for the correct formation of mRNP export platform to coordinate the proper mRNP remodeling process at cytoplasmic end of nuclear channel ^38,40–42^.

### Inner and outer Nup205 play different roles in CR

For the question mark shaped density attached to the Y complex arms in CR, previous studies speculated that it might be Nup205 or Nup188 ^14,20,34^. Recently, by taking a closer view with higher resolution structure of *X. laevis* NPC, the densities were attributed to Nup205 in both CR and NR ^27,28^. Here, according to the isotropic density map of CR subunit and AlphaFold2 predictions, we modeled full-length Nup205 structure toward the question mark shaped densities attached to both inner and outer Y complexes, named as inner and outer Nup205 (Fig. 4A-D). As control, we also tried the modeling of Nup188 according to the same local density. Similar as our previous report for NR ^28^, the structure of Nup205 fits both inner and outer densities much better than Nup188 (Fig. 4E). The most obvious difference is the tower helix in the middle domain of Nup205, which is missing in Nup188.

**Figure 4.**
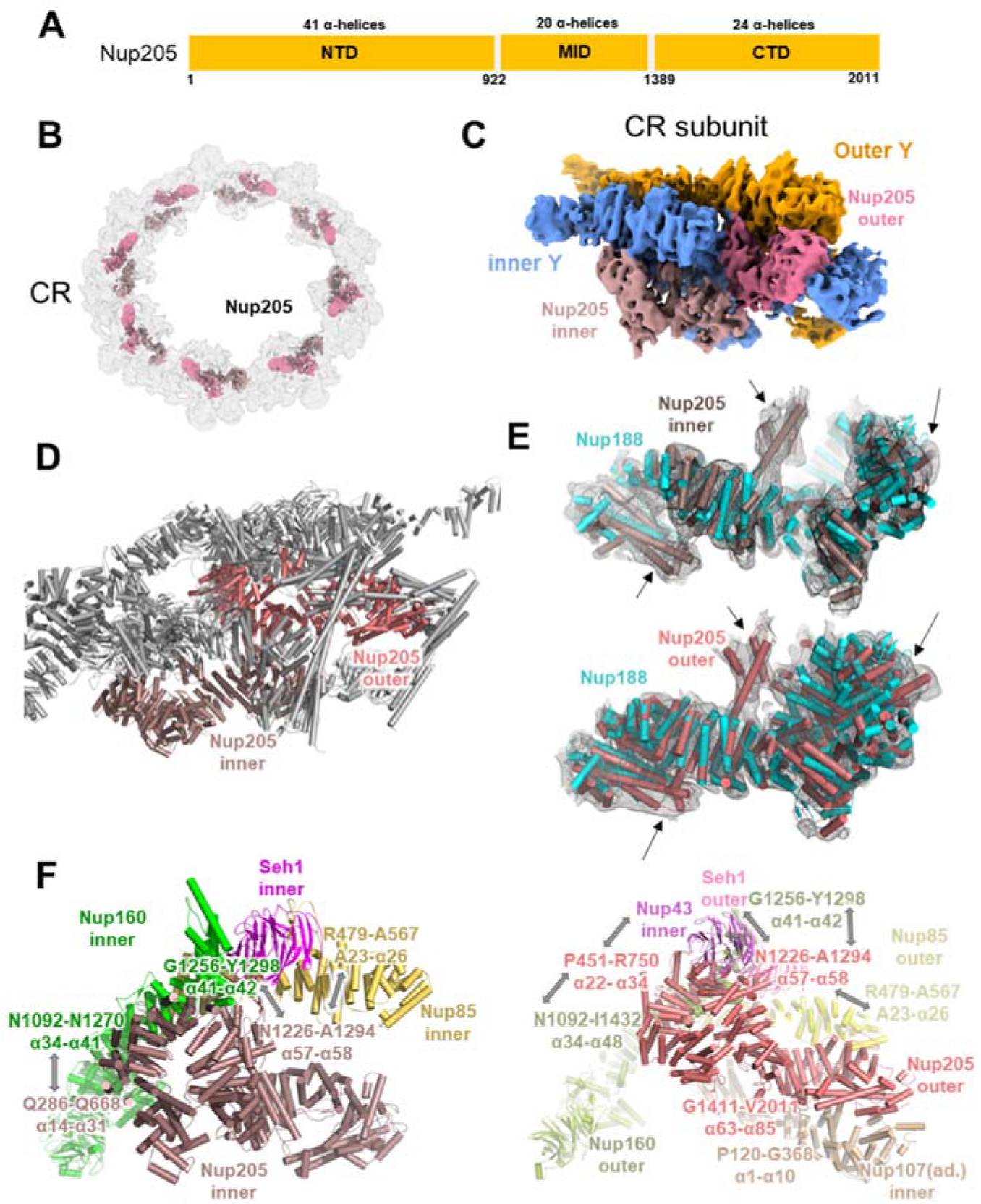
The structures and interaction details of inner and outer Nup205 in CR subunit. (**A**) Domain assignment of Nup205. (**B**) Location of 16 copies Nup205 in CR, showing relative positions of inner and outer Nup205, Nup205 within each subunit were colored differently. (**C**) Location of 2 copies Nup205 in CR subunit, inner Nup205 was colored in salmon, outer Nup205 was in pale violet red. (**D**) Model of inner and outer Nup205 attached to CR subunit. (**E**) Model overlay of Nup205 and Nup188 onto local densities, showing major differences. (**F**) Interaction sites of inner and outer Nup205 to surrounding Nups. a.d., adjacent subunit.

According to the improved model of inner and outer Nup205, we found that they have quite different interaction features with surrounding Nups. For inner Nup205, it NTD (α14 to α31 helices) contacts with inner Nup160 (α41 to α42 helices), while its tower helix region (α57 to α58 helices) interacts with inner Nup85, inner Seh1 and inner Nup160 (Fig. 4F). For outer Nup205, its NTD connects to inner Nup43 and outer Nup160, the tower helix region binds to outer Nup85, outer Seh1 and outer Nup160, the CTD interact with Nup107 of adjacent CR subunit (Fig. 4F). It showed that inner and outer Nup205 help to stabilize the two Y complexes formation and head-to-tail fashion in CR.

### Nup93 connects Y complex stems in CR subunit

Recently, we revealed that in NR subunit of *X. laevis* NPC, Nup93 acts as a bridge to connect two Y complexes in the stem (Fig. 5A) ^28^. But the corresponding location in CR subunit has always been regarded as the density for Nup358, so whether there is a similar Nup93 bridge in CR as is still unclear to date. According to the improved cryo-EM map of CR subunit, we found that when the density corresponding to the five copies of Nup358 is removed from CR subunit, an unassigned density like ‘bridge domain’ emerges between two Y complex stems (Fig. 5B-D). Using AlphaFold2’s prediction as the starting reference, we modelled 31 α-helices of Nup93 into this local density (Extended Data Fig. 3H). Besides its interactions with the five Nup358 proteins as shown above, it also connects multiple Y complex Nups. In Nup93 NTD, its α1 to α17 helices (P173 to P542) connects to inner Sec13 and inner Nup96’s α13 to α15 helices (S595 to G669). In its CTD, the α23 to α31 helices (P636 to N820) connects to regions near outer Nup107’s finger helix α28 to α35 (L621 to Q861) (Fig. 5D). Therefore, this α-solenoid domain of Nup93 CR participated in the formation of two Y complexes assembly through its interaction with both Y complexes’ stems in CR.

**Figure 5.**
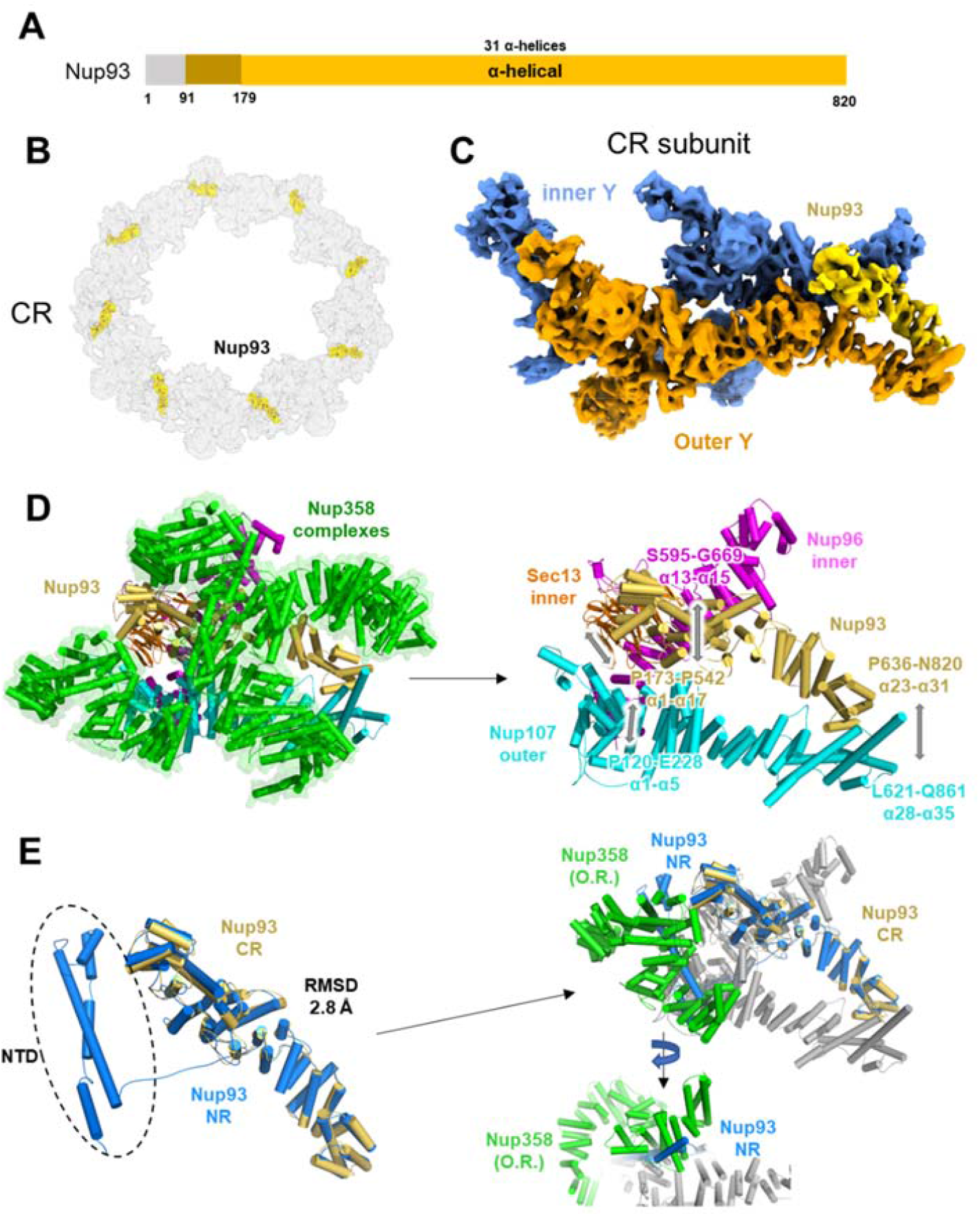
Nup93 acts as a bridge to connect the stems of the inner and outer Y complexes in CR. (**A**) Domain assignment of modeled part of Nup93. (**B**) Location of 8 copies Nup93 in CR, colored in gold. (**C**) Location of Nup93 in CR subunit. (**D**) Interaction sites of Nup93 with surrounding Nups. (**E**) Model comparison between Nup93 in CR and NR, showing spatial conflicts between outer right Nup358 with NTD of Nup93 from NR.

When comparing the Nup93 models from NR and CR, we found that the CR Nup93 have 5 helices missing in the NTD (Fig. 5E), so it seems shorter than NR Nup93. In the superposition of these two models, their main bodies share high structural similarity with a RMSD value of 2.8 Å (Fig. 5E). The location for the missing helices of Nup93 NTD in CR is fully occupied by outer-right Nup358. Therefore, it might be possible that Nup358 binds to Y complex more tightly than Nup93 NTD and occupies its interface, then the density corresponding to Nup93 NTD is missing in CR subunit (Fig. 5E).

## Discussion

For a long time, there are two major obstacles in solving the detailed structure of NPC using SPA method. One is the anisotropic resolution caused by preferred orientation problem, the other is lack of accurate full-length structure of all Nups in NPC. Here, we combined “side-view” particles and “tilt-view” particles to overcome the insufficient Fourier space sampling problem and used AlphaFold2 to predict all Nups’ structures. Based on the isotropic reconstruction map (Extended Data Fig. 2, 7) and the highly accurate predicted models (Extended Data Fig. 4), we managed to build a most comprehensive and accurate pseudoatomic model of the NPC CR to date. The multiple novel structural features in this model represent a great advance in understanding the assembly of NPCs.

The localization of Nup358 onto both stems of Y complexes was initially established by knockdown experiments of NPC in Hela cell ^14^. Then multiple studies proved that Nup358 is vital for double Y complexes arrangement in CR subunit, since those species lacking Nup358 has only one Y complex left in each CR subunit ^14,16^. For the first time, we found that there should be at least five copies of Nup358 warped around the stem regions of Y complexes in each CR subunit, and each Nup358 forms extensive but different interactions with surrounding Nups. This result agrees well with previous reports for the importance of Nup358. It’s not easy to understand why NPC needs so many copies of Nup358 in the CR region. One possible reason may be related to the large cytoplasmic filament attached to CR. Its high dynamic requires the stable connection to the CR base, so it needs multiple copies of Nup358 proteins for itself to be firmly anchored on the CR.

The exportation of mRNAs from nucleoplasm to cytoplasm relies on the correct modeling and remodeling of mRNPs, while remodeling of mRNPs on cytoplasmic side of NPC requires participation of Gle1, Nup42, Nup214 and DEAD-box helicase ^41,43^. The mRNP remodeling platform was proposed to project from cytoplasmic ring onto the nuclear channel to ensure the efficient mRNP remodeling process. Here, we found that there are two Nup214 complexes in each CR subunit of *X. laevis* NPC, both pointing to the nuclear channel to facilitate correct formation of mRNP export platform. However, in yeast, the bases of this mRNP remodeling platform were reported to be a Nup82 holo-complex or a P shaped architecture ^17,38^. The structural difference of the Nup214 complex in different species may be related to the different copy number of Y complexes in CR. In yeast, the NPC has only one Y complex in each CR subunit, so it cannot support the rake-shaped structure of Nup214 (Nup159) complex that found in this study, since parallel formation of Nup214 complexes requires the connection to both inner and outer Nup85 simultaneously. Then in yeast, Nup159 needs to form a different confirmation, such as the holo-complex dimer, to anchor onto the CR region. Moreover, the two copies of Nup214 complexes in parallel seem to provide a much denser arrangement of FG-repeat domains inside the nuclear channel, which may lead to a more efficient mRNP exporting process in vertebrates than yeast.

According to our much improved models of CR in this study and NR in previous report ^28^, there are 2 copies of Nup205 in CR subunit and 1 copies in NR subunit. In the reported low-resolution model of IR, there should be 4 Nup188 or Nup205 in each IR subunit ^20,34^. Meanwhile, according to stoichiometry data for Nups reported previously, the total amount of Nup205 in NPC is roughly the twice of Nup188 ^7^. Hence, it’s almost impossible that there is only Nup205 or only Nup188 in IR, and it should be the combination of Nup205 and Nup188, but the exact result requires high-resolution model of IR in the future.

In summary, we solved the cryo-EM map of the *X. laevis* NPC CR in an isotropic resolution around 8 Å and obtained a more accurate and complete model at secondary structure level (Supplementary Video 1). The revealed new structural details advanced our understanding toward the detailed organization and assembly of vertebrate NPC.

## Methods

### Sample preparation

The sample preparation of African clawed toad X. laevis oocyte nucleus envelope has been described in details previously ^28^. Briefly, ovaries were removed from narcotized mature female *X. laevis*, and stage VI oocytes were isolated, and NE were applied onto the grid in ice-cold HEPES buffer (83 mM KCl, 17 mM NaCl, 10 mM HEPES, pH 7.5). Before plunge freezing, the sample on the grid were cross-linked with 0.15% glutaraldehyde for 10 min on ice. After cross-linking process, the grid was blotted and vitrified by plunge freezing into liquid ethane by Vitrobot Mark IV (Thermo Fisher Scientific, USA) at 100% humidity, all grids were stored in liquid nitrogen before imaging.

The animal experiments were performed in the Laboratory Animal Center of Peking University in accordance with the National Institutes of Health Guide for the Care and Use of Laboratory Animals and according to guidelines approved by the Institutional Animal Care and Use Committee at Peking University.

### Cryo-EM data acquisition

The data acquisition collection strategy of this study was basically the same with our previous reports ^28^. Briefly, after screening in Talos Arctica 200 KV TEM (Thermo Fisher Scientific, USA), the good grids were mounted into Titan Krios 300 KV TEM (Thermo Fisher Scientific, USA) for imaging. 8745 images were collected at a nominal magnification of 64,000X, resulted in a calibrated physical pixel size at the specimen level of 2.24 Å. For images at tilting angle at 0/30/45/60 degrees, the total dose was set to be 100 or 120/60/80/100 e-/Å2, the movies were recorded on a 0.5 s per frame base, and the exposure time of these collected data set were set to be 28.5 or 34.5/21.5/41 or 28.5/36 or 35 s (Extended Data Table 1). All movies were recorded by a Gatan K2 Summit DDD detector (Gatan Company, USA) under super resolution mode, equipped with a post column GIF Quantum filter, whose slit width was set to be 20 eV. SerialEM with in-house scripts was used for data collection with the defocus value set between 1.0 to 4.0 μm ^44,45^.

### SPA image processing

The image processing workflow was basically the same with our previous reports ^28^. Briefly, motion correction along with dose weighting were performed by MotionCor2 ^46^. Particle picking was done by using RELION ver3.0 prior to CTF estimation ^47^. Only particles with apparent feature as NPC were kept for further processing. CTF estimation was done using Gctf or goCTF or Warp on per-particle basis ^48–50^.

The Image alignment processing of CR were done using RELION 3.0 unless stated specifically ^47^. Prior to alignment of CR, we docked the previously reported model of the CR from human NPC (PDB entry 5A9Q) into full NPC map, and segmented surrounding density using Chimera. The segmented part was used to generate a mask solely covering CR part in our map ^14,51^. Then refinement of CR on binned 4 level using this CR mask was done. The initial reference was generated by low pass filtered the reported human NPC structure to 60 Å, and C8 symmetry was applied during refinement. The refinement of CR on binned 4 level reached a final resolution of 29 Å. Then, using refined orientations and shifts, CR particle at binned 2 level were extracted with a box size of 400 pixels. After extraction, reconstruction was done for the extracted particles to generate a mask solely covering CR region. Using similar strategy as at binned 4 level of CR particles, refinement was done and reached a final resolution of 23 Å.

Then, the alignment was done on subunit level, as no significant gain of resolution would be achieved for whole CR by decreasing binning levels. The relative coordinate of CR subunit to CR box center was determined using Chimera, then we used a modified version of block-based reconstruction script to generate a RELION star file containing orientations and updated defocus values of each subunit ^51,52^. Then, we extract the subunit particles and first ran a reconstruct job to make sure everything was right. The model of PDB 5A9Q was used to generate a mask solely covering regions corresponding to one asymmetric unit, and refinement using this mask was done to reach a resolution of 10.7 Å for CR subunit. Extraction of binned 1 particles was done using refined shifts and orientations from refinement of binned 2 particles, with a box size of 320 pixels. Like what was done for binned 2 CR subunit particles, first a reconstruction was done to obtain an initial reference and a mask solely covering one subunit was generated. Refinement at this stage reached a resolution of 9.8 Å. Next, we ran Bayesian Polishing of all particles. The output star file was separated into multiple files, each containing particles corresponding to individual stage tilting angles. Then 3D classification was done for these individual tilts, using the refined map as reference. After classification, all particles corresponding to the best class in different jobs were selected and merged, then an auto refinement was done for the classified particles and reached a resolution of 8.8 Å. Then the output star file and corresponding map were subjected to CryoSPARC for final refinement using its local refinement tool and resulted in a final resolution of 8.7 Å ^53^. Similarly, a mask covering the most rigid part of CR subunit was created, also subjected to CryoSPARC for local refinement using the same particles data set and reached a final resolution of 8 Å for CR core region ^53^. A similar strategy was applied to CR Nup358 region and reached a final resolution of 8.9 Å. The validation of map and model quality of this research was done using 3D-FSC and Phenix ^30,54^(Extended Data Fig.2).

### Modeling of NPC CR

The full version of AlphaFold2 was installed as instructed with all database downloaded ^29^. All the structures of NPC CR Nups from *X. laevis* or *X. tropicalis* (Nup160 and Nup96), were predicted by AlphaFOLD2 using the recommended parameters. Briefly, the value of Max_template_hits was set to 20, Relax_energy_tolerance was set to 2.39, Relax_stiffness was set to 10, Relax_max_outer_iterations was set to 20. For each Nup, a total of 5 relaxed structures were predicted, and the prediction with the highest confidence was selected as the starting model for the next refinement.

Then we performed stepwise MDFF simulations to refine each Nup model according to the corresponding local density in CR subunit. A timestep of 1 fs was used throughout the simulation. Langevin dynamics were adopted at a temperature of 310 K. The equilibration step for energy minimization was performed on the initial model for 1000 steps before the refinement run. The refinement runs were performed for 3000 ps, which corresponds to 3,000,000 simulation steps, and the gridForceScale values were gradually increased fromb 0.3 to 0.7 during the refinement. All simulations were performed using CHARMM36m forcefields ^55^. Electrostatic calculations were treated with particle mesh Ewald (PME). A cutoff of 12 Å was chosen for short-range van der Waals interactions. NAMD ^56^ was used as the MD engine throughout all simulations.

All the CR components were assembled in COOT ^57^ for manually adjustment according to the overall density map of CR subunit. Then the whole model of CR subunit was refined using PHENIX.real_space_refine ^58^. Data collection statistics and refinement statistics are given in Extended Data Table 1. All figures in this study were generated by PyMol, Chimera and ChimeraX ^51,59^.

## Data Availability

The Electron Microscopy Database (EMD) accession codes of the CR subunit region, CR core region and CR Nup358 region are EMD-32056, EMD-32060, EMD-32061, respectively. The Protein Data Bank (PDB) accession code of the model of the CR subunit is 7VOP.

## ACKNOWLEDGEMENTS

We thank all other members of the Fei Sun and Chuanmao Zhang laboratories for their help. We would also like to thank the Center for Biological Imaging (CBI), Institute of Biophysics, Chinese Academy of Science for cryo-EM work, and Boling Zhu, Xujing Li, Gang Ji, Jiashu Xu and Guoliang Yin for their help with cryo-EM data collection; the Facilities Cores at National Center for Protein Sciences and the cryo-EM and TEM platforms at the College of Life Sciences of Peking University for cryo-electron microscopy and TEM; and Ning Gao, Zhenxi Guo, Guopeng Wang, Yingchun Hu, Xia Pei and Bo Shao for their help with cryo-EM and TEM experiments.

This work was equally supported by grants from Ministry of Science and Technology of China (2017YFA0504700 to FS and 2016YFA0500201 to CMZ), the Strategic Priority Research Program of the Chinese Academy of Sciences (XDB 37040102 to FS), and National Natural Science Foundation of China (31830020 to FS, 31520103906 to CMZ). This work was also supported by grants from the National Science Fund for Distinguished Young Scholars (31925026 to FS), National Natural Science Foundation of China (31430051 to CMZ) and National Key Research and Development Program of China (2016YFA0100501 to CMZ and 2018YFA0901102 to YZ).

## AUTHOR CONTRIBUTIONS

F. S. and C. Z. conceived the project and designed the experiments. L. T., H. R. and X. H. performed cryo-EM experiments. L. T. and Y. Z. performed cryo-EM data processing. H. R. and L. T. participated in the preparation and screening of cryo-EM samples. Y. Z. performed the modeling and simulation-based refinement. L. T. and Y. Z. analyzed the data and wrote the manuscript, which was substantially revised by F. S. and C. Z.

