## Supplemental Data for "8 Å structure of the cytoplasmic ring of the *Xenopus laevis* nuclear pore complex solved by cryo-EM and AI"

### Extended Data figures and tables

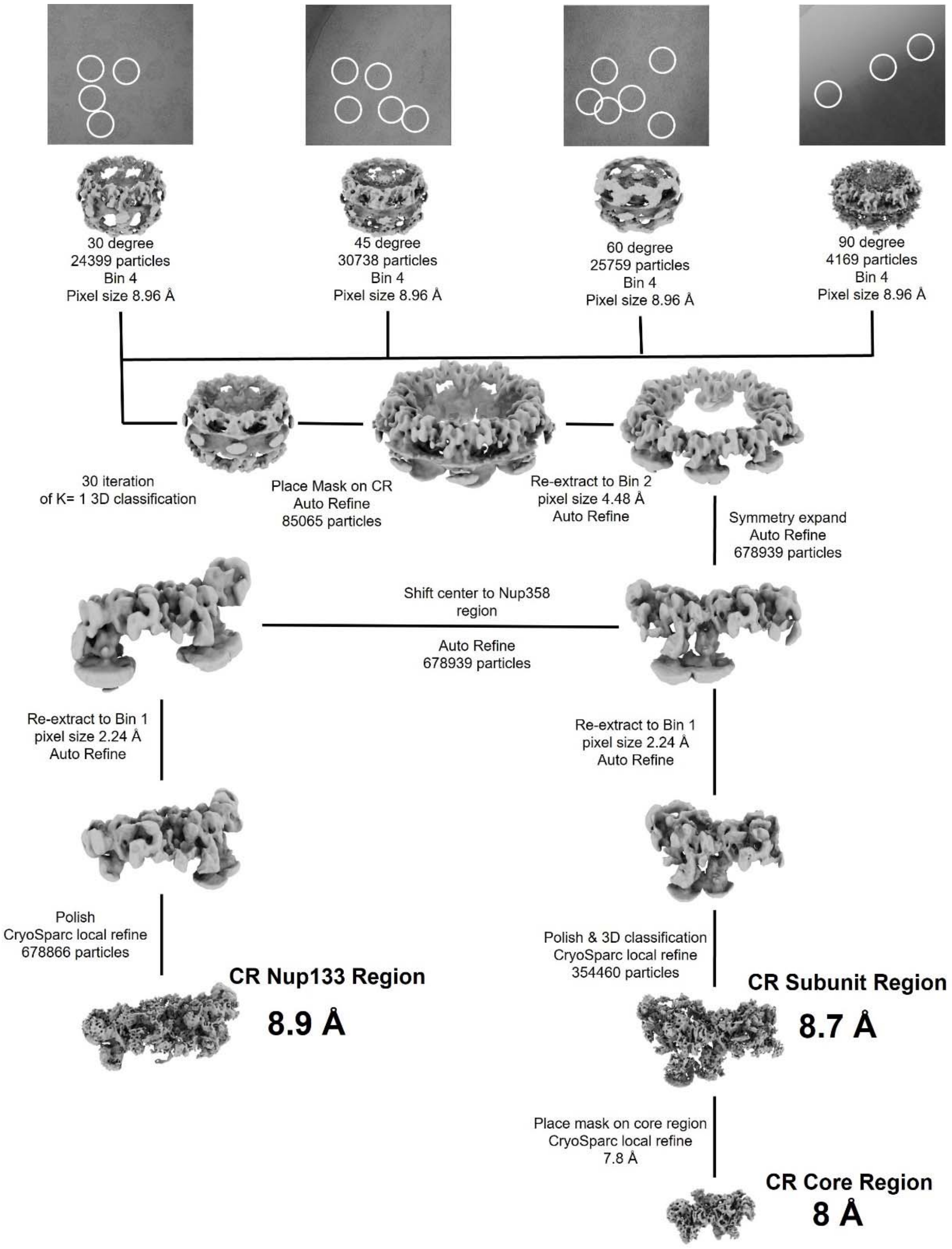


**Extended Data Figure 1. Data processing workflow.**

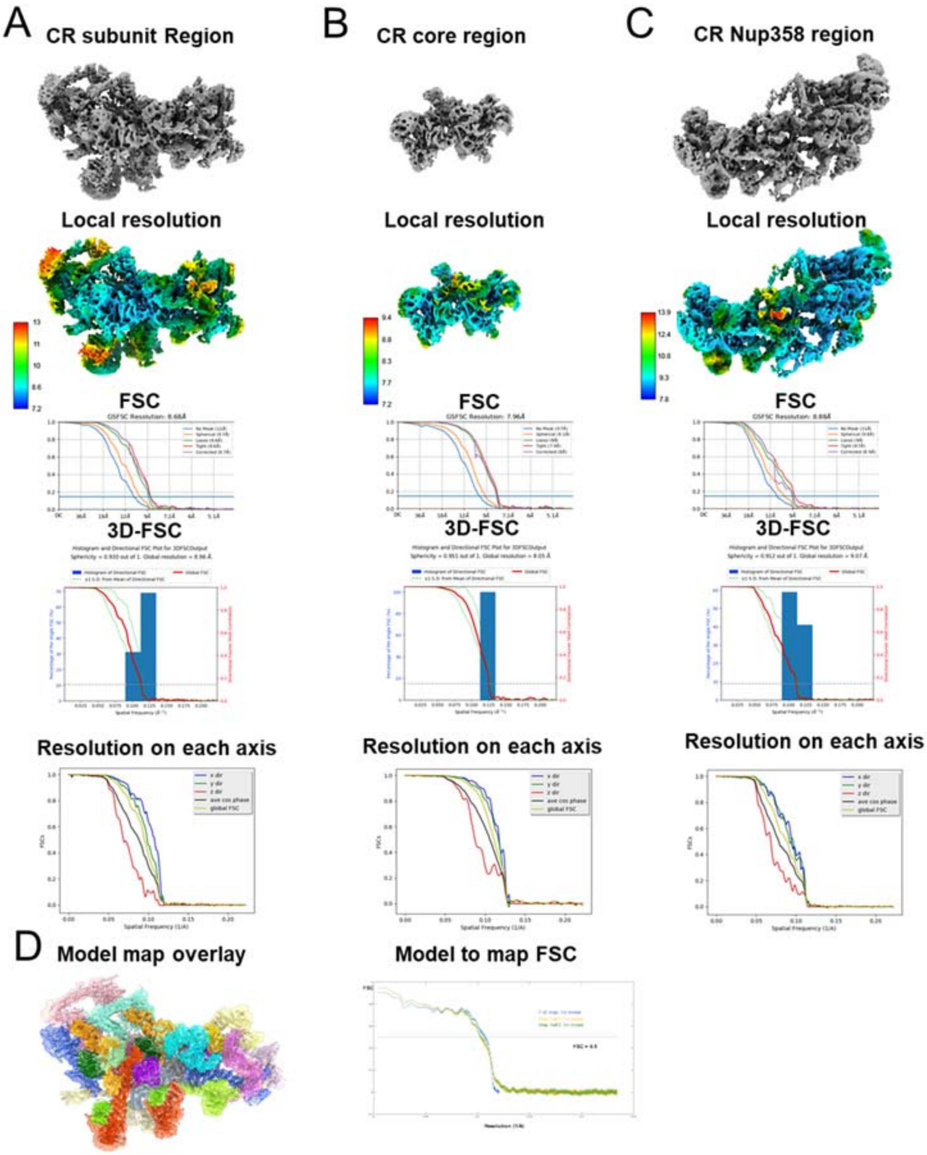


**Extended Data Figure 2. Assessments of cryo-EM maps. (A, B, C)** Map display, local resolution estimation, FSC, 3D-FSC and directional FSC of the CR subunit region, CR core region and CR Nup358 region. **(D)** Map model overlay of the CR subunit, and map versus model FSC estimation.

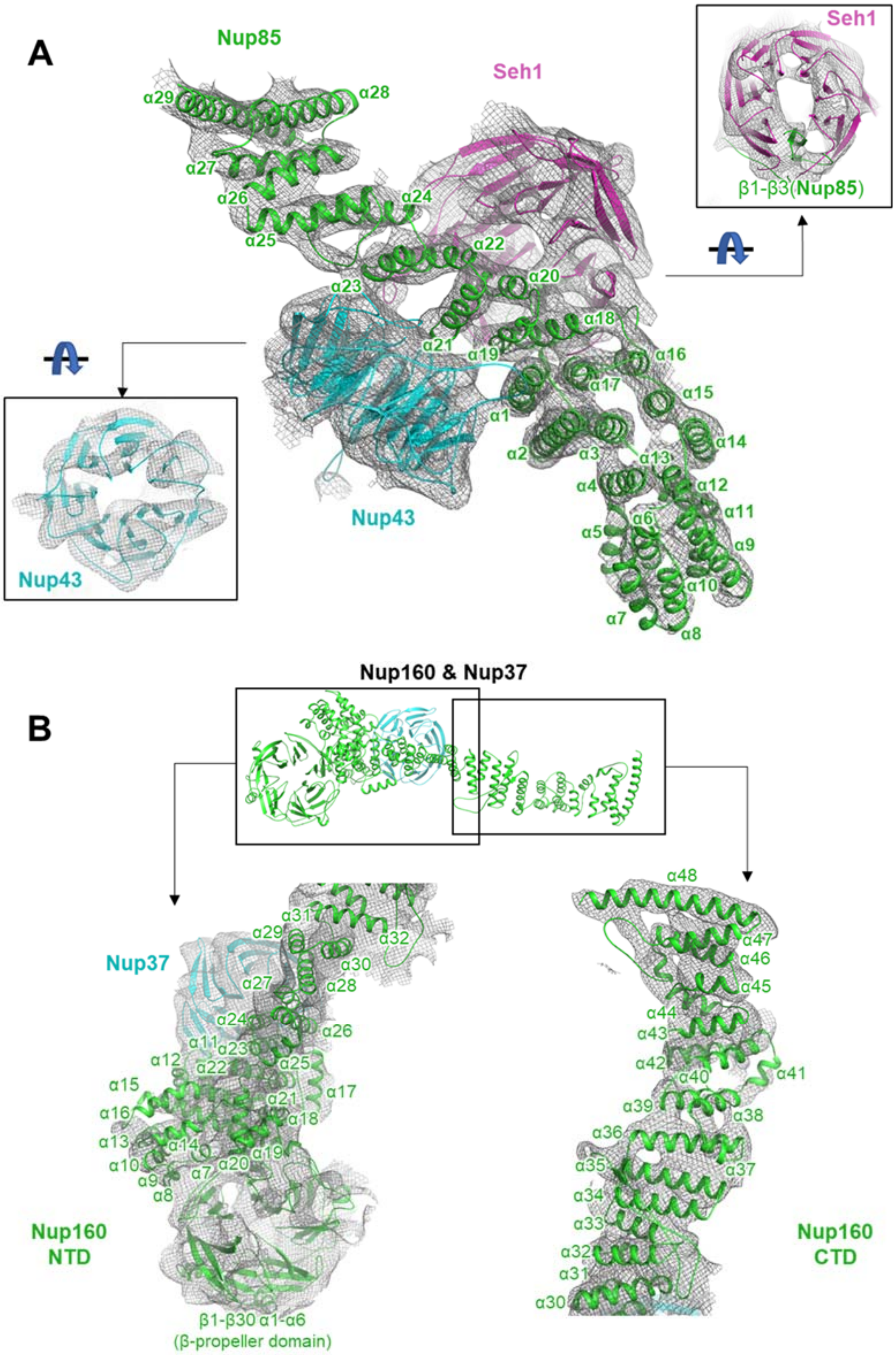

A
Nup85
Seh1
α29
α28
α27
α26
α24
α25
α22
α20
α23
α18
α21
α19
α16
α17
α15
α1
α14
α2
α3
α13
α4
α12
α11
α5
α6
α9
α10
α7
α8
Nup43
Nup43
Seh1
β1-β3(Nup85)
B
Nup160 & Nup37
Nup37
α31
α29
α32
α30
α27
α28
α24
α26
α11
α23
α12
α22
α25
α15
α17
α16
α21
α18
α13
α14
α10
α7
α20
α19
α9
α8
Nup160
NTD
β1-β30 α1-α6
(β-propeller domain)
α48
α47
α46
α45
α44
α43
α42
α41
α40
α39
α38
α36
α35
α37
α34
α33
α32
α31
α30
Nup160
CTD

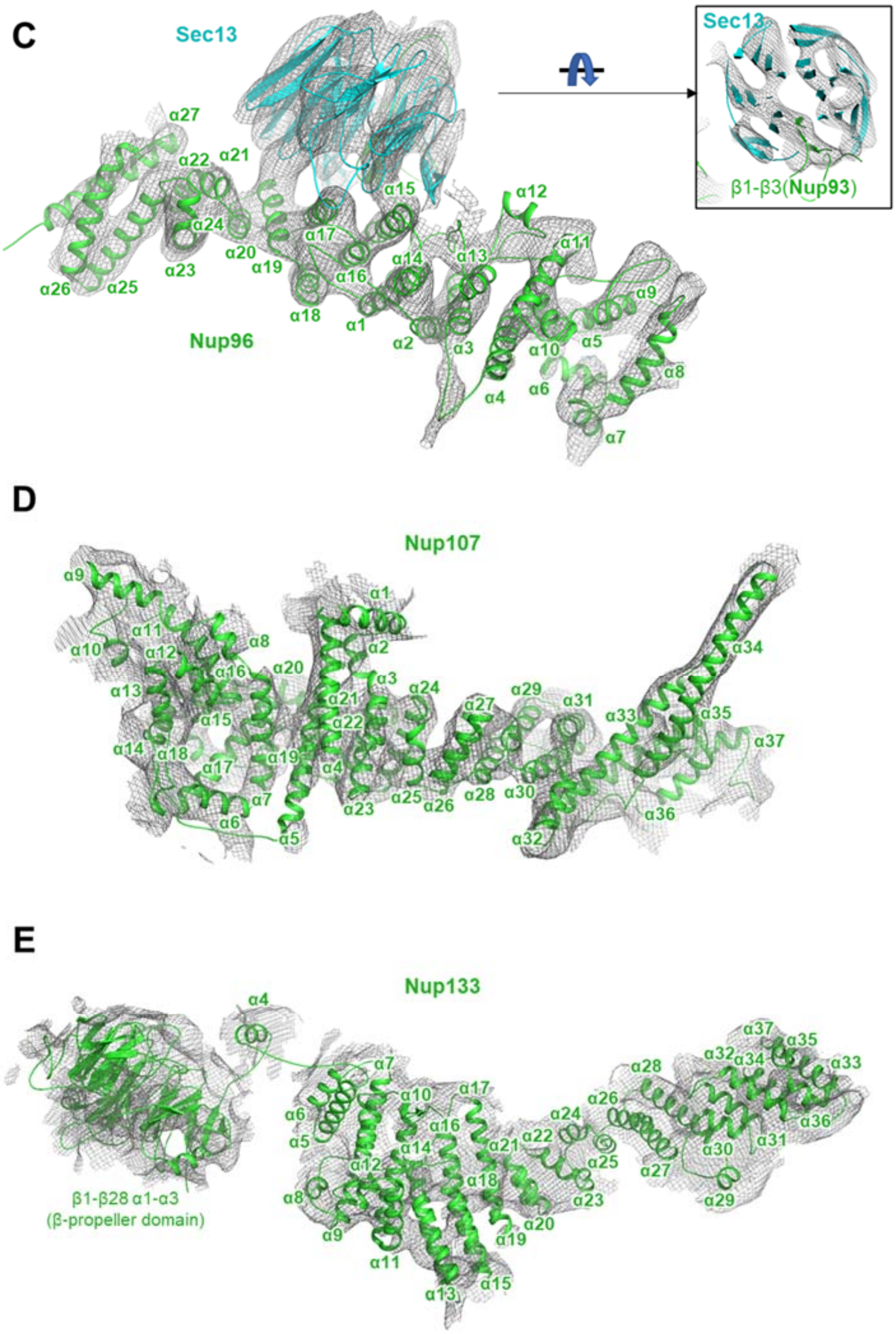
C
Sec13
Sec13
β1-β3(Nup93)
Nup96
D
Nup107
E
Nup133
β1-β28 α1-α3
(β-propeller domain)

**F**

**Inner Nup205**

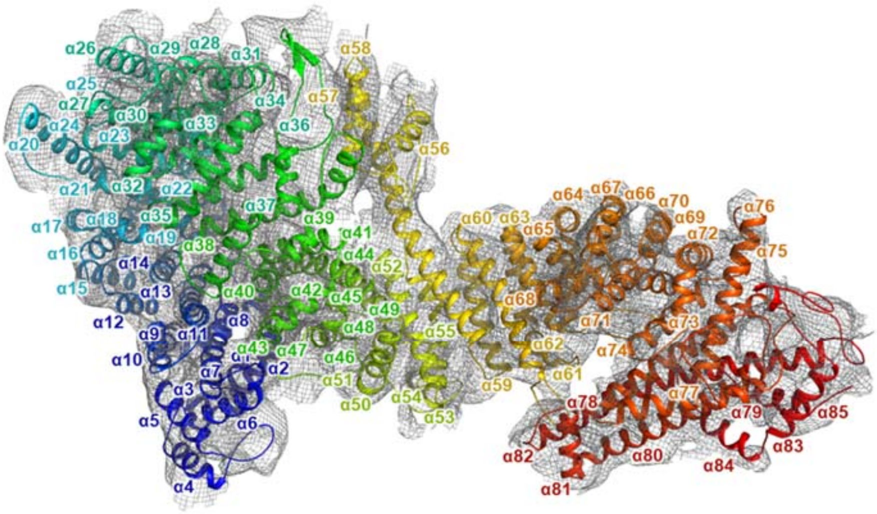


**G**

**Outer Nup205**

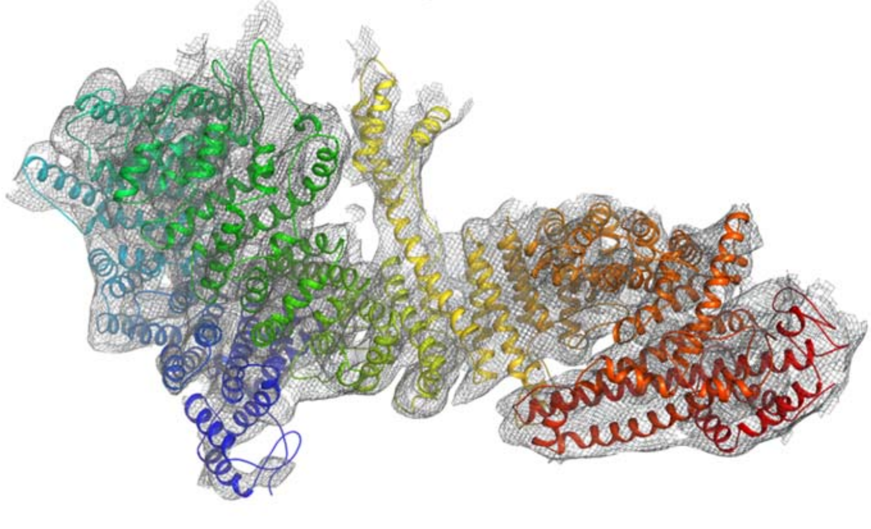

**H**

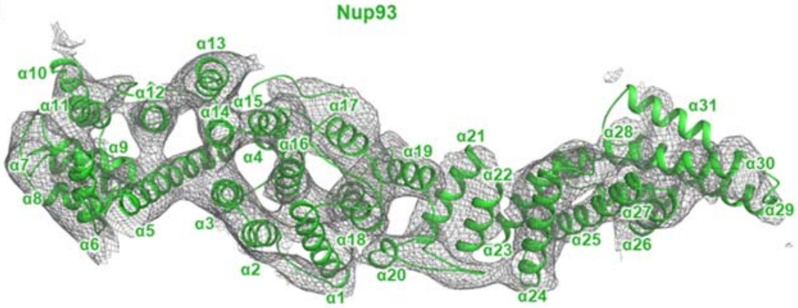
Nup93
α1
α2
α3
α4
α5
α6
α7
α8
α9
α10
α11
α12
α13
α14
α15
α16
α17
α18
α19
α20
α21
α22
α23
α24
α25
α26
α27
α28
α29
α30
α31

**I**

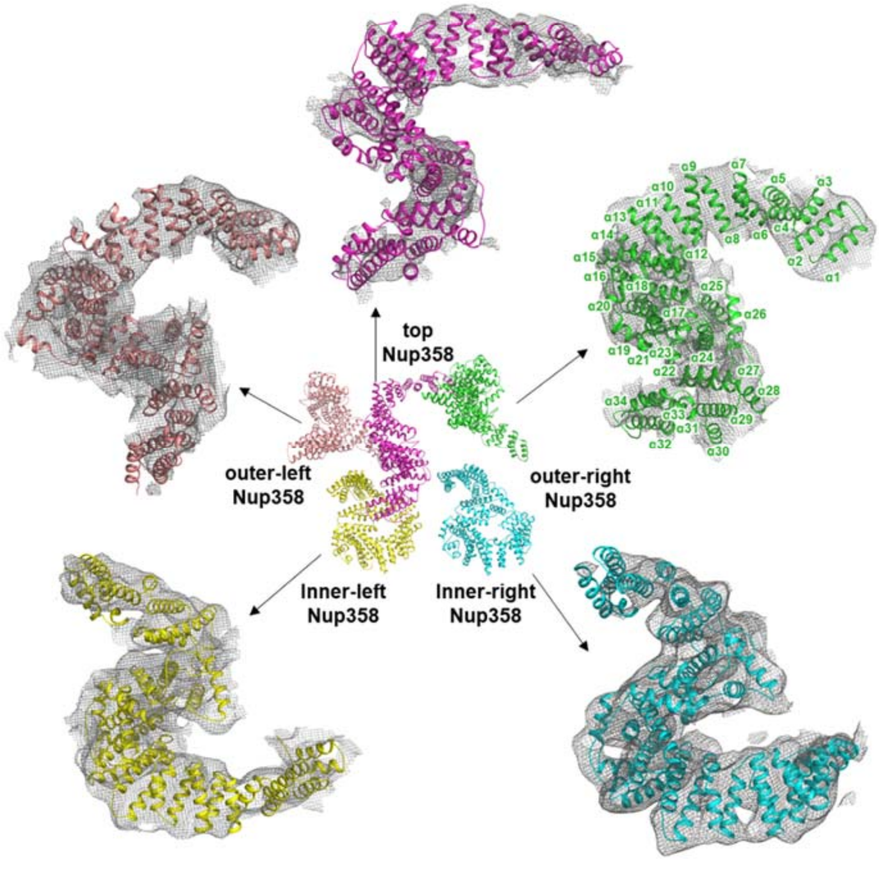
top
Nup358
outer-left
Nup358
outer-right
Nup358
Inner-left
Nup358
Inner-right
Nup358

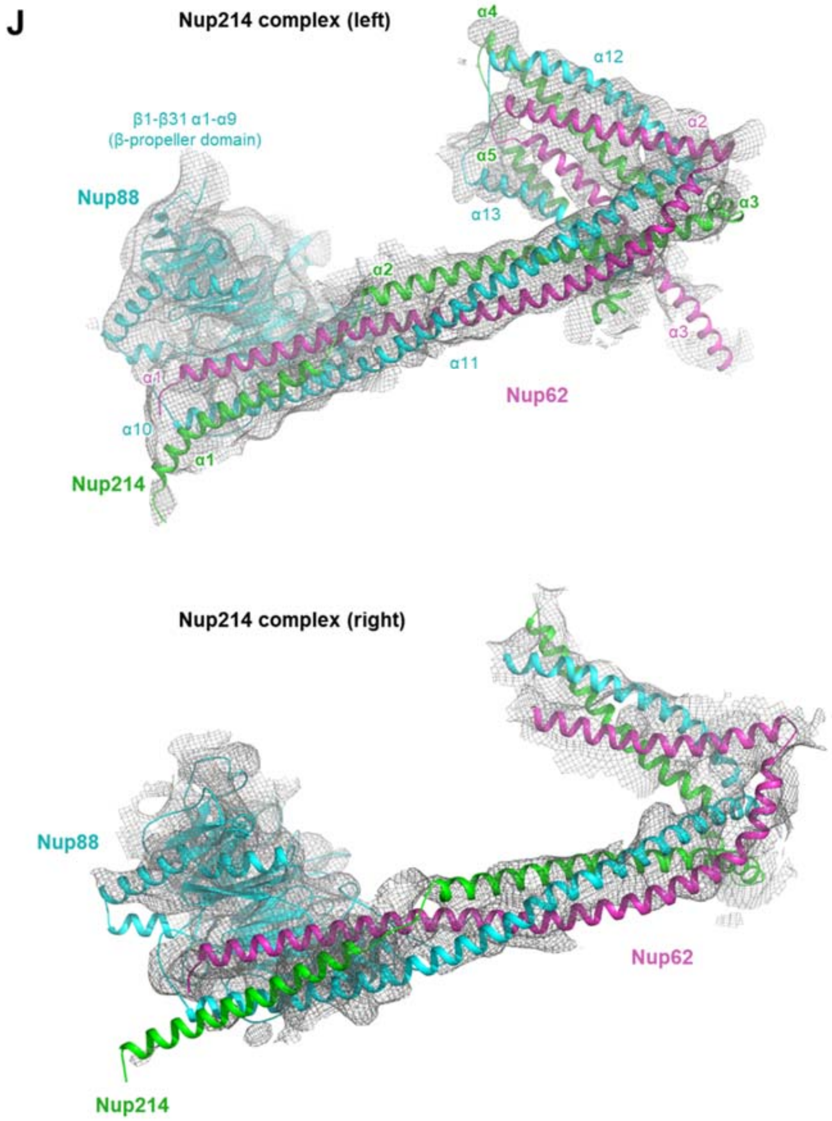


**Extended Data Figure 3. Model building quality and secondary structure assignment of the CR subunit. (A-J)** Model building quality of Nup85, Seh1, Nup43, Nup160, Nup37, Sec13, Nup96, Nup107, Nup133, Nup205, Nup93, Nup358 and Nup214 complexes.

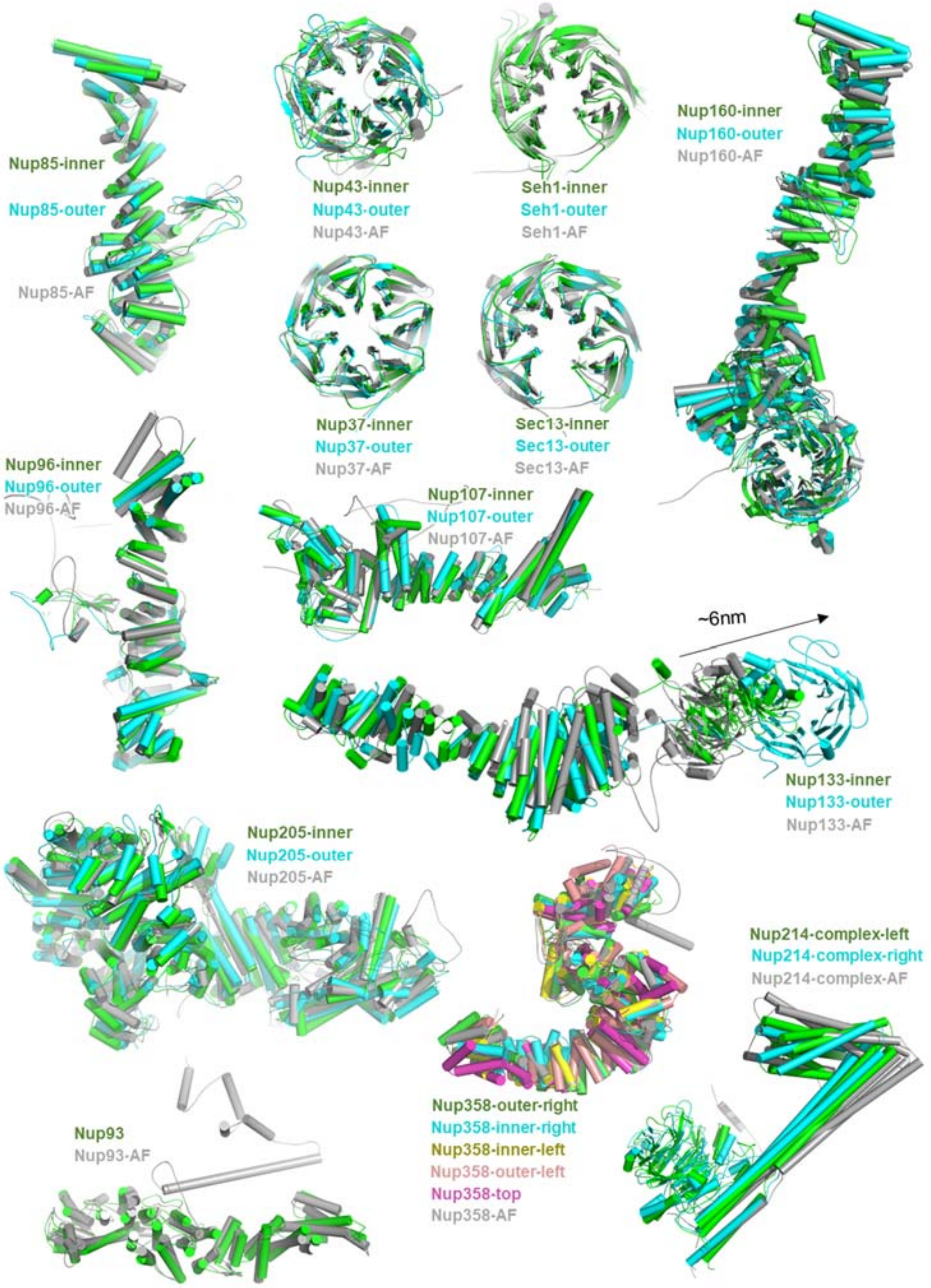


**Extended Data Figure 4. Comparisons of the refined structure and predicted structure by AlphaFold2 (AF) of CR Nups from the *X. laevis* NPC.**

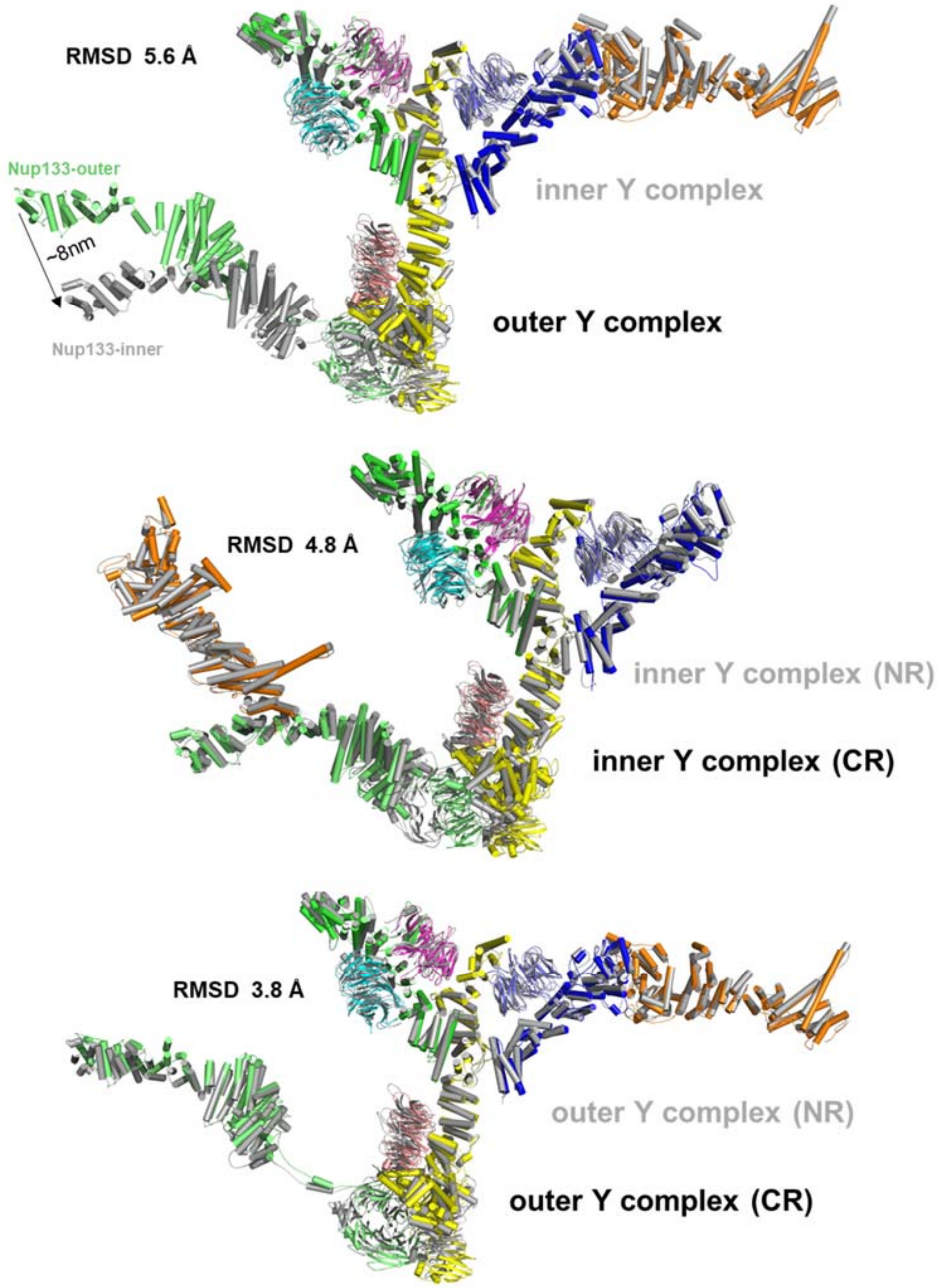


**Extended Data Figure 5. Structural comparations of Y-complexes in NR & CR from *X. laevis* NPC.**

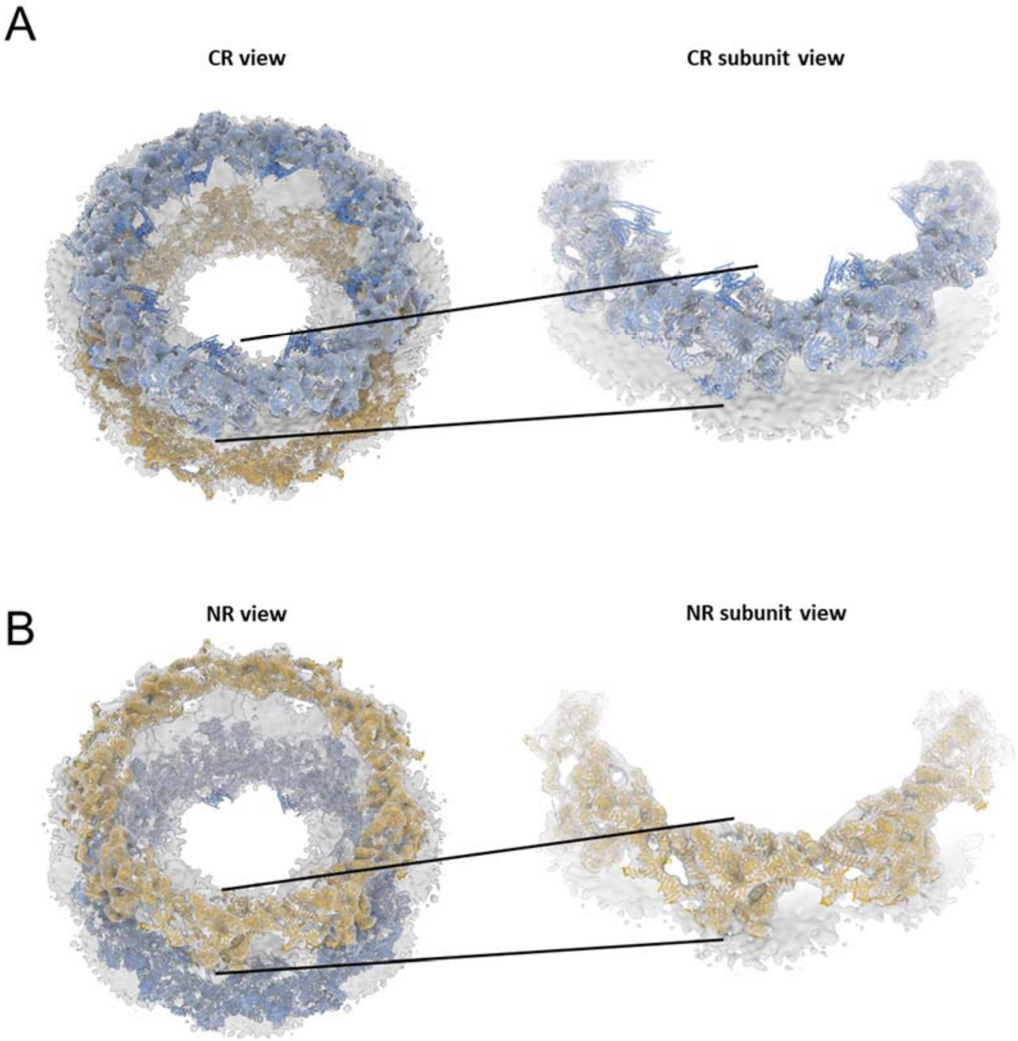


**Extended Data Figure 6. NR and CR models of *X. laevis* NPC fitted in human NPC map (EMD-3103).**

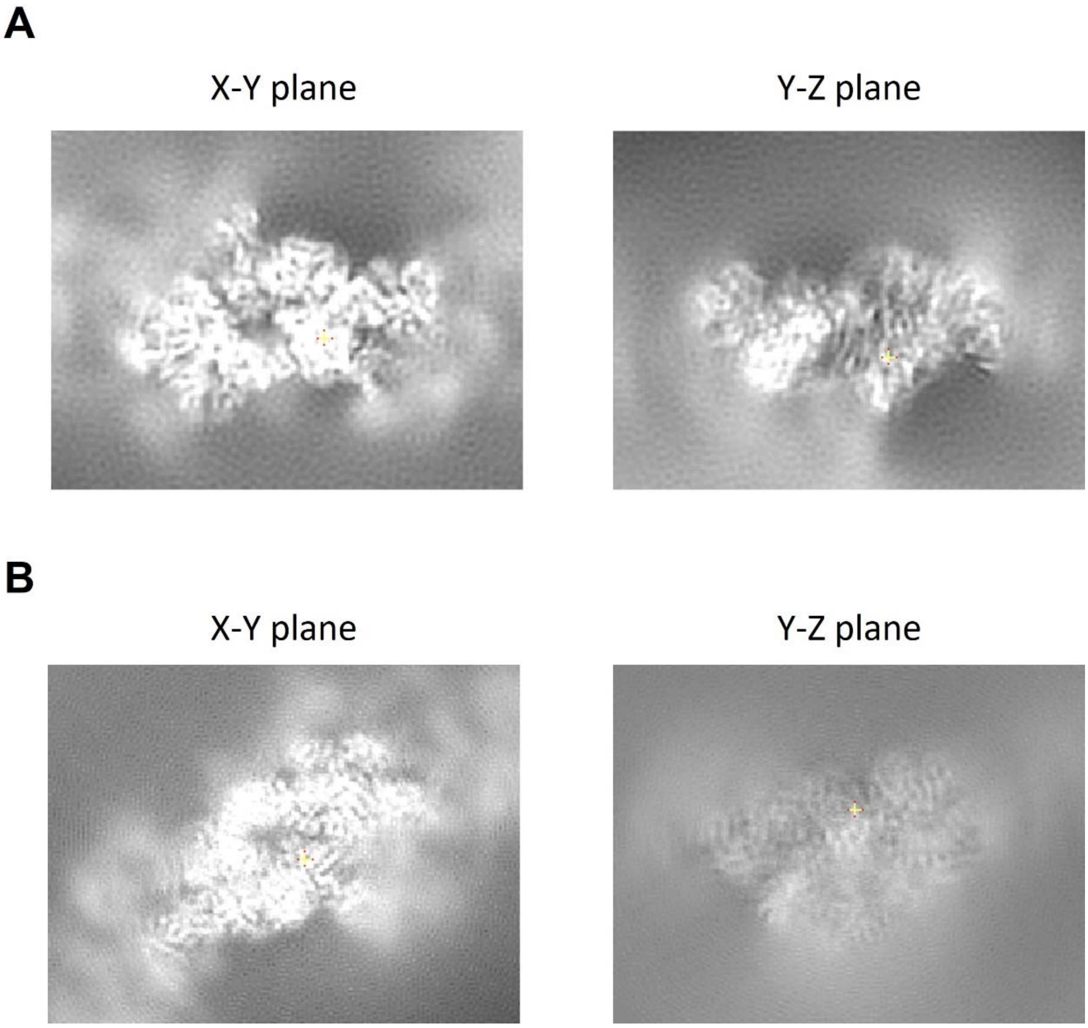


**Extended Data Figure 7. Comparison of slices viewing from different axis of (A) CR core region of our map and (B) previous reported *X. laevis* CR core map (EMD-0909).**

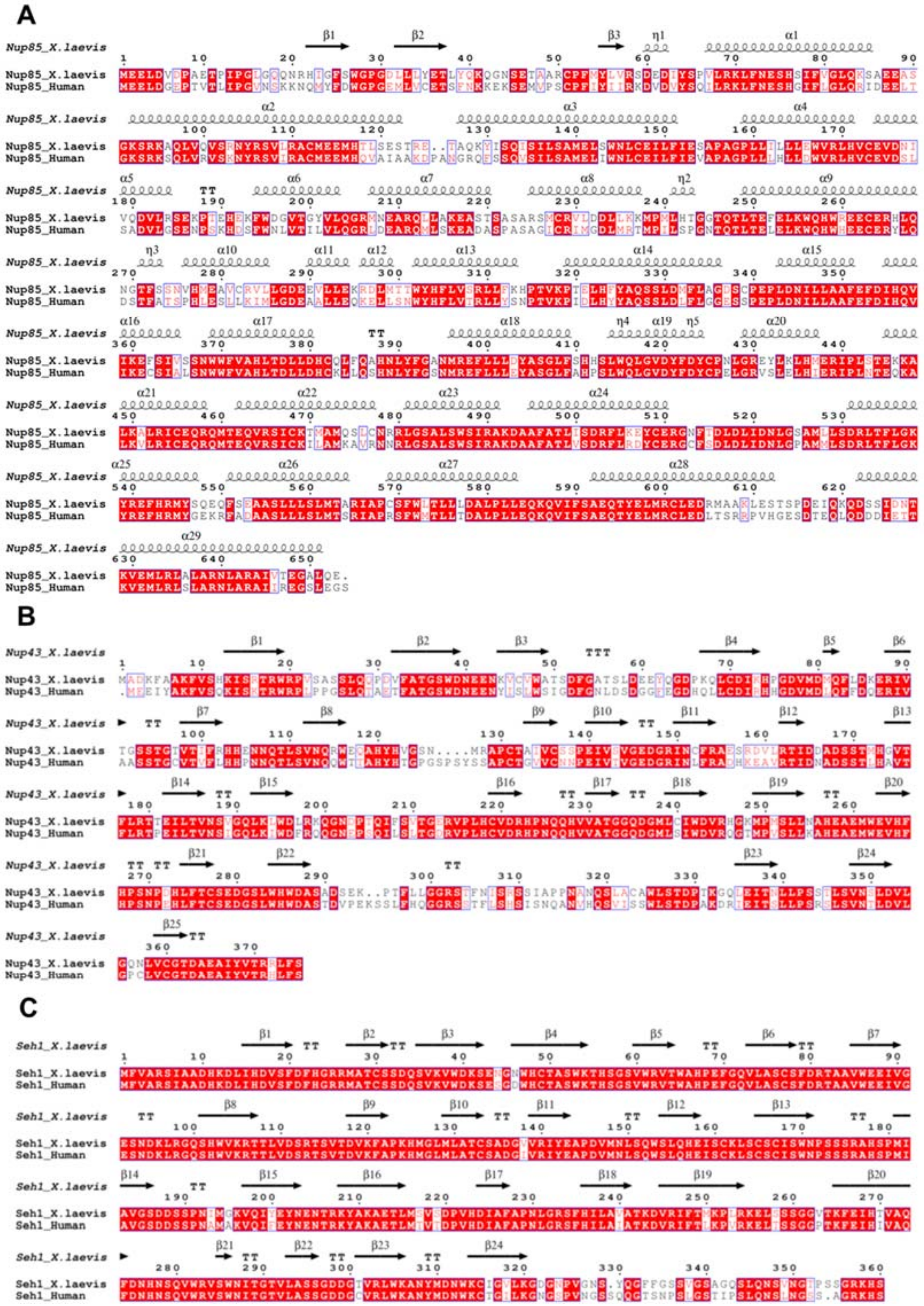
A
Nup85_X.laevis
Nup85_Human
B
Nup43_X.laevis
Nup43_Human
C
Seh1_X.laevis
Seh1_Human

**D**

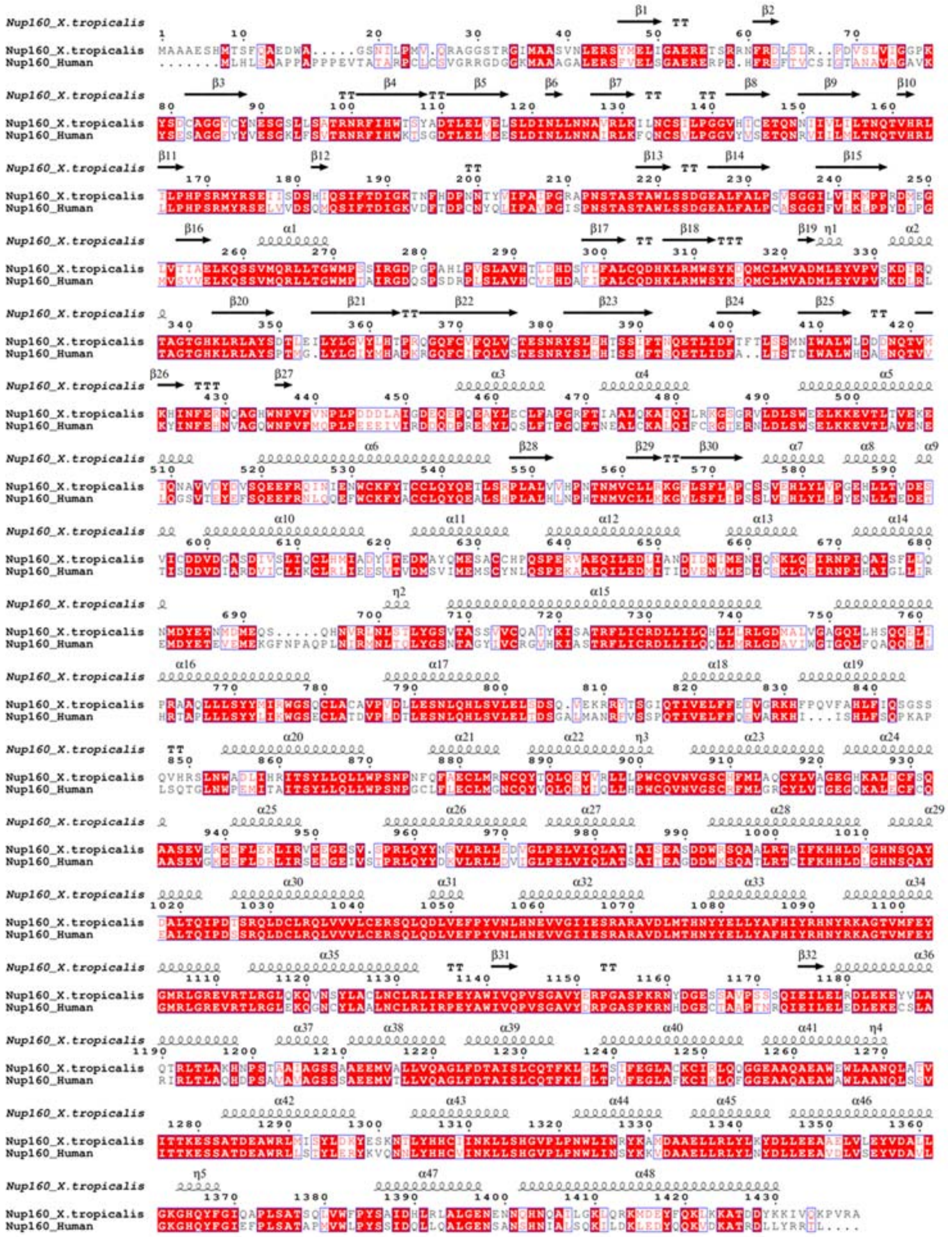

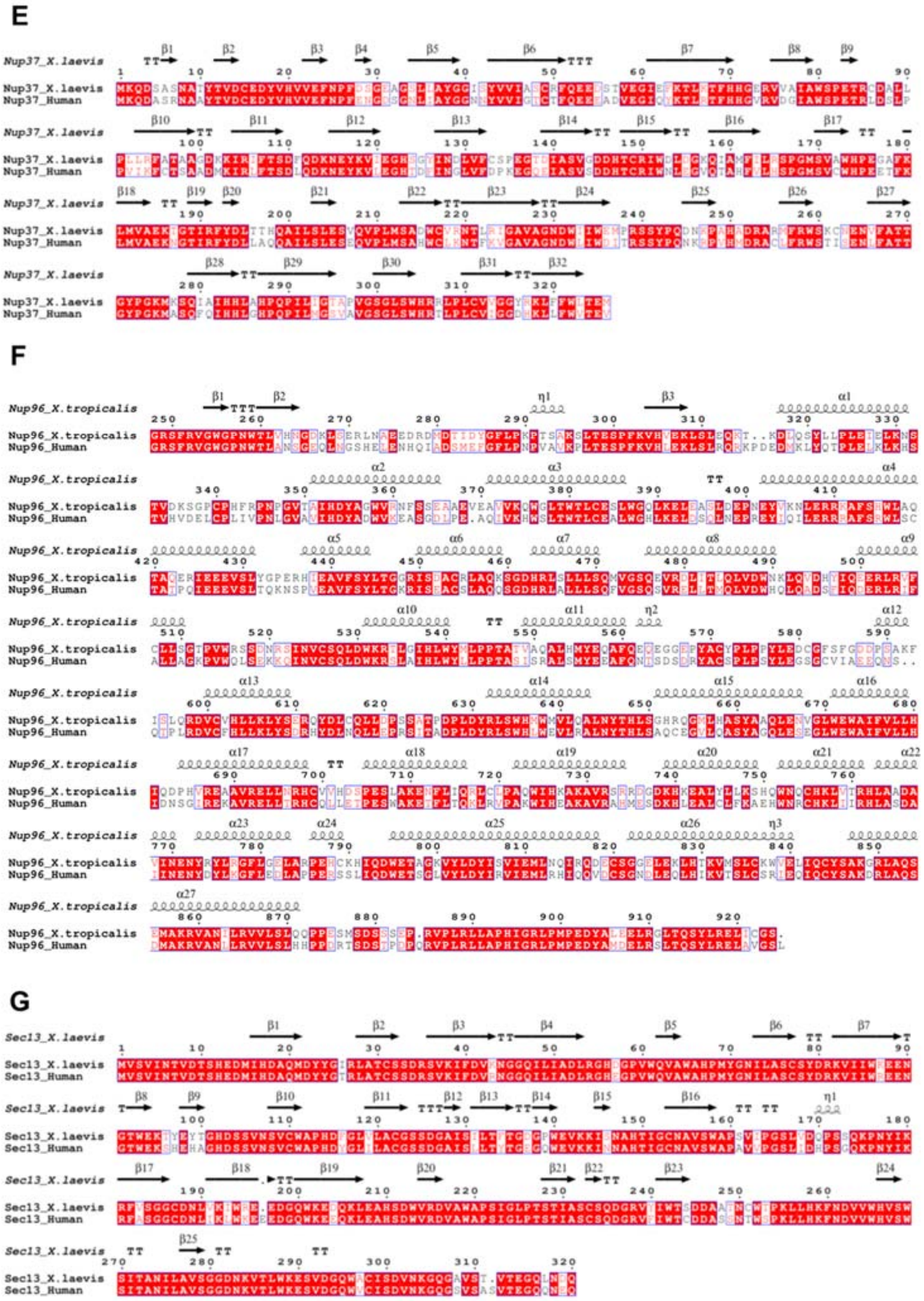
E
Nup37_X.laevis
Nup37_Human
F
Nup96_X.tropicalis
Nup96_Human
G
Sec13_X.laevis
Sec13_Human

H

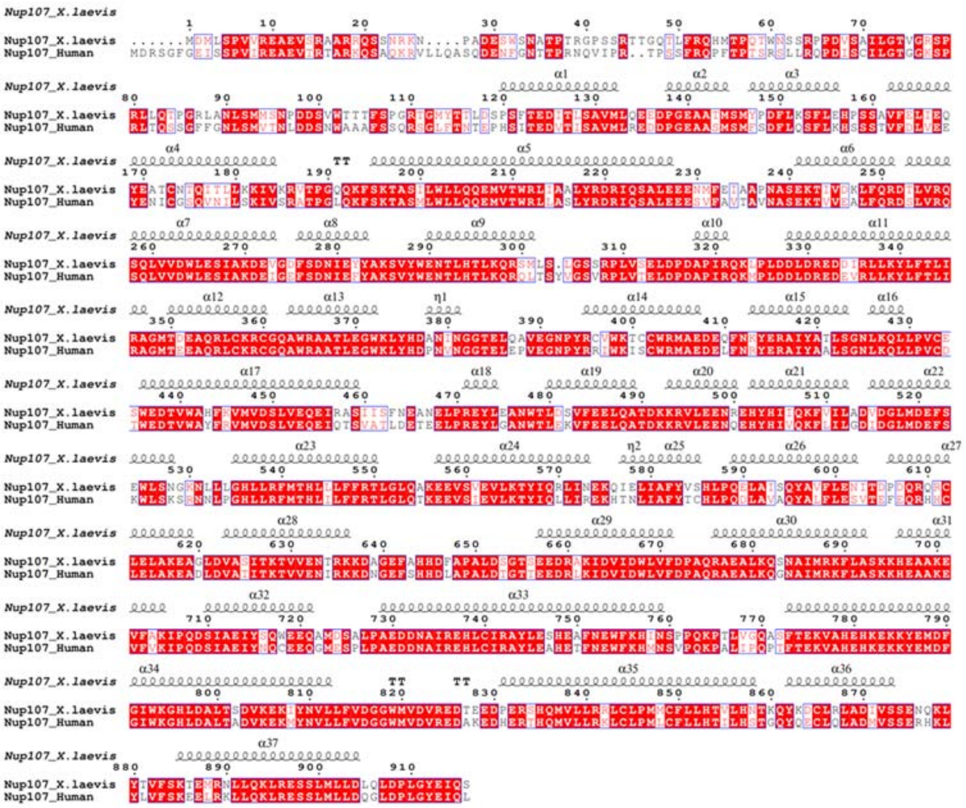

I

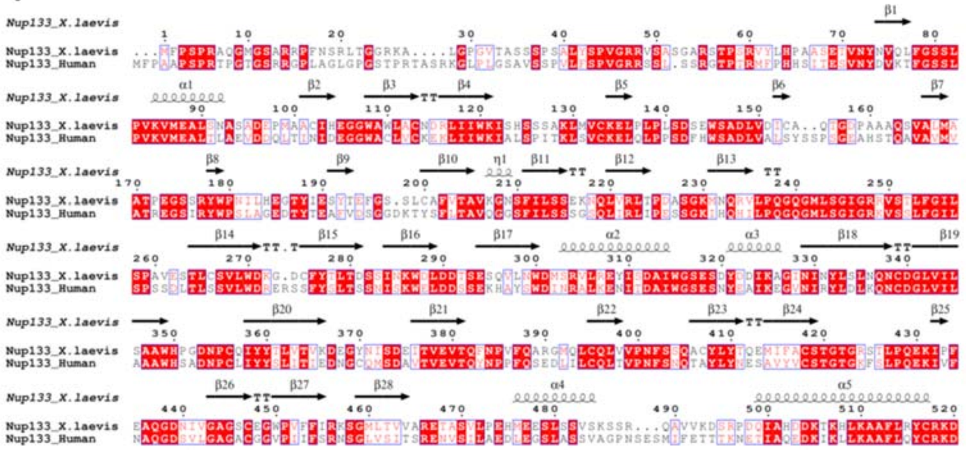

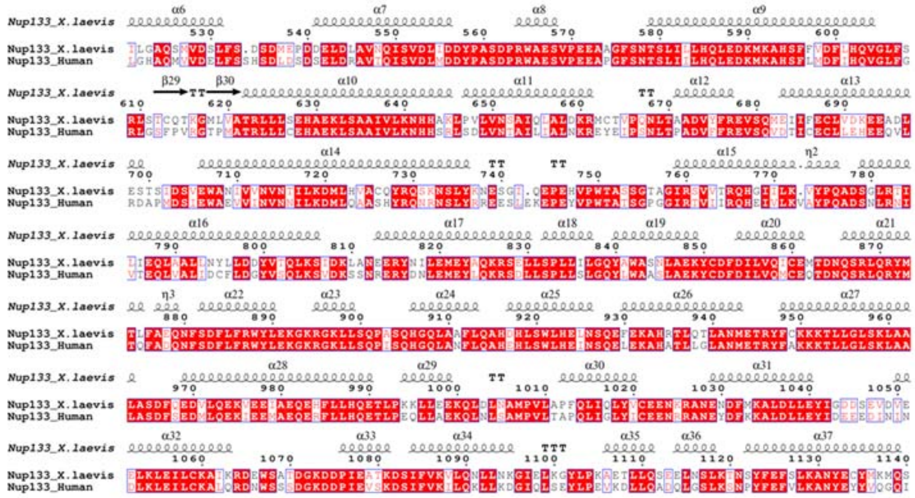

J

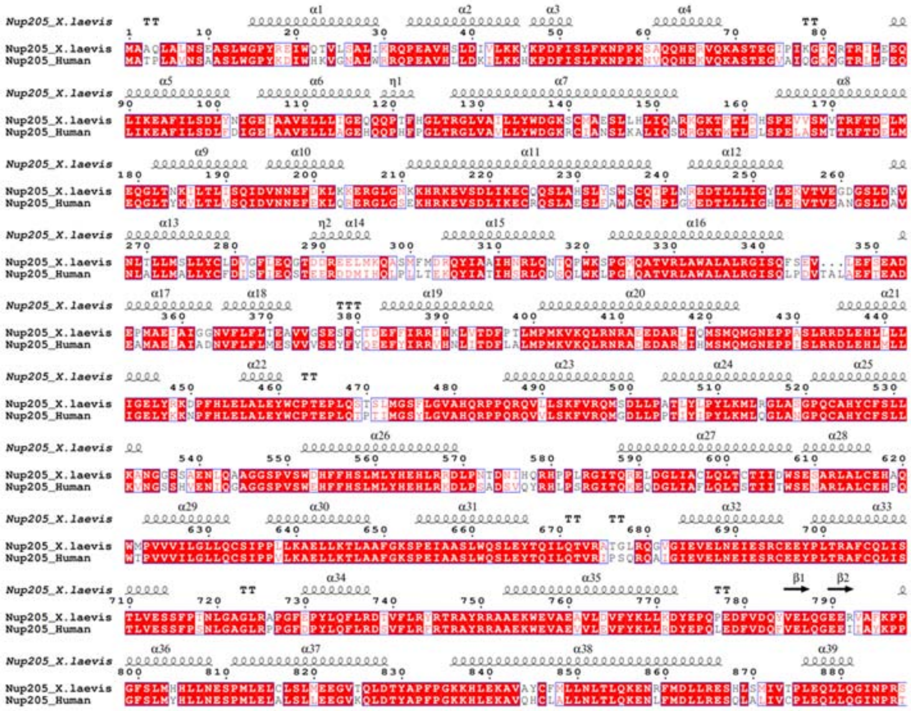

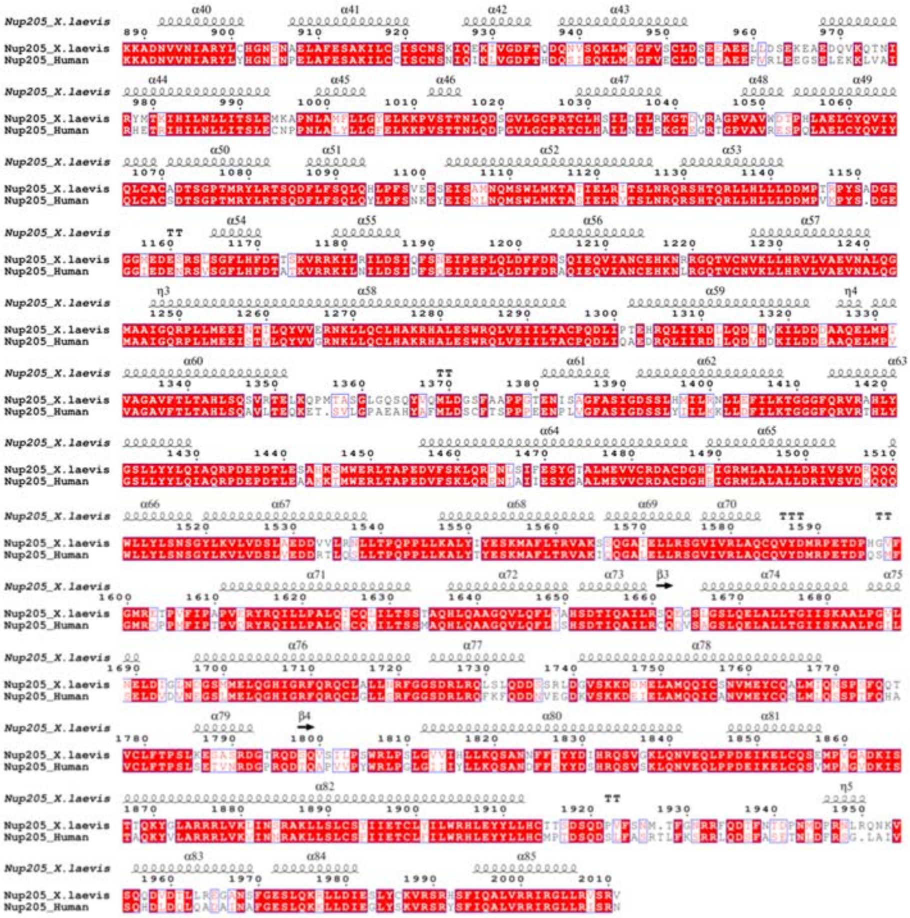

K

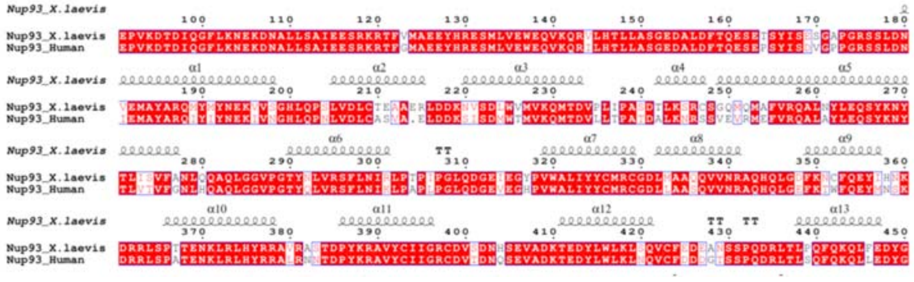

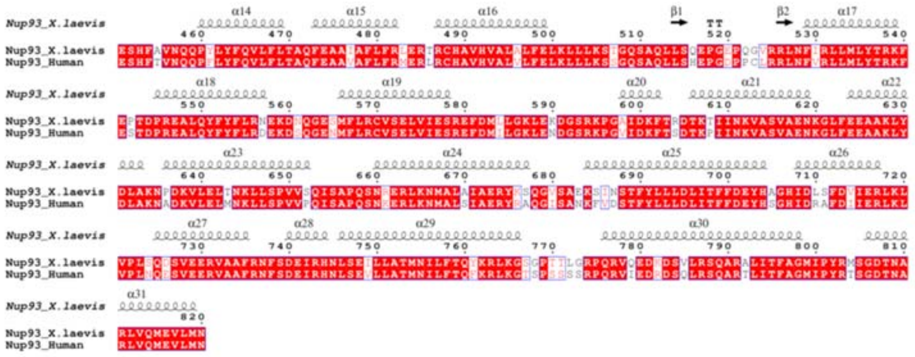


L

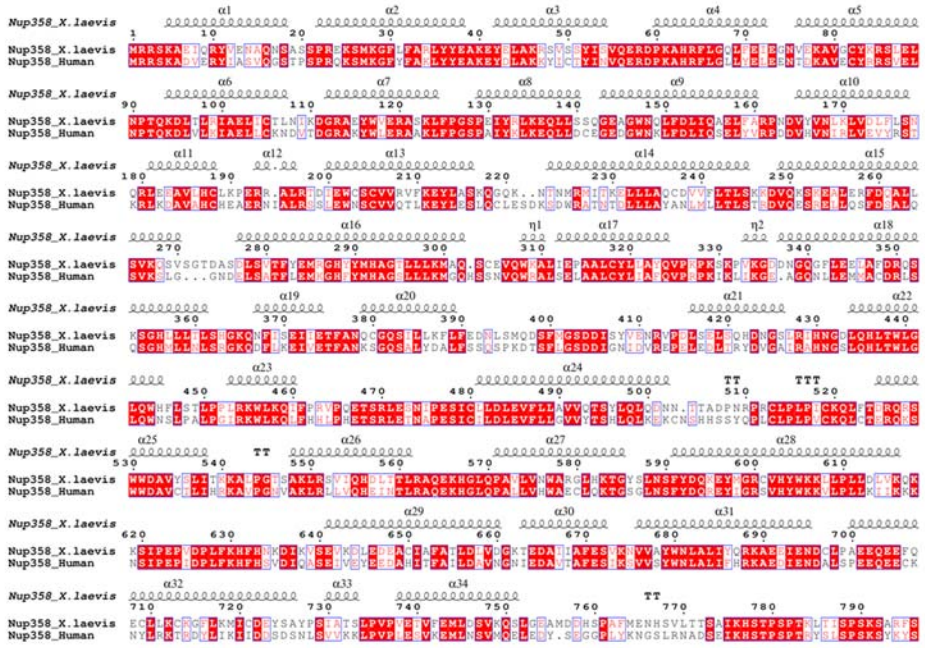

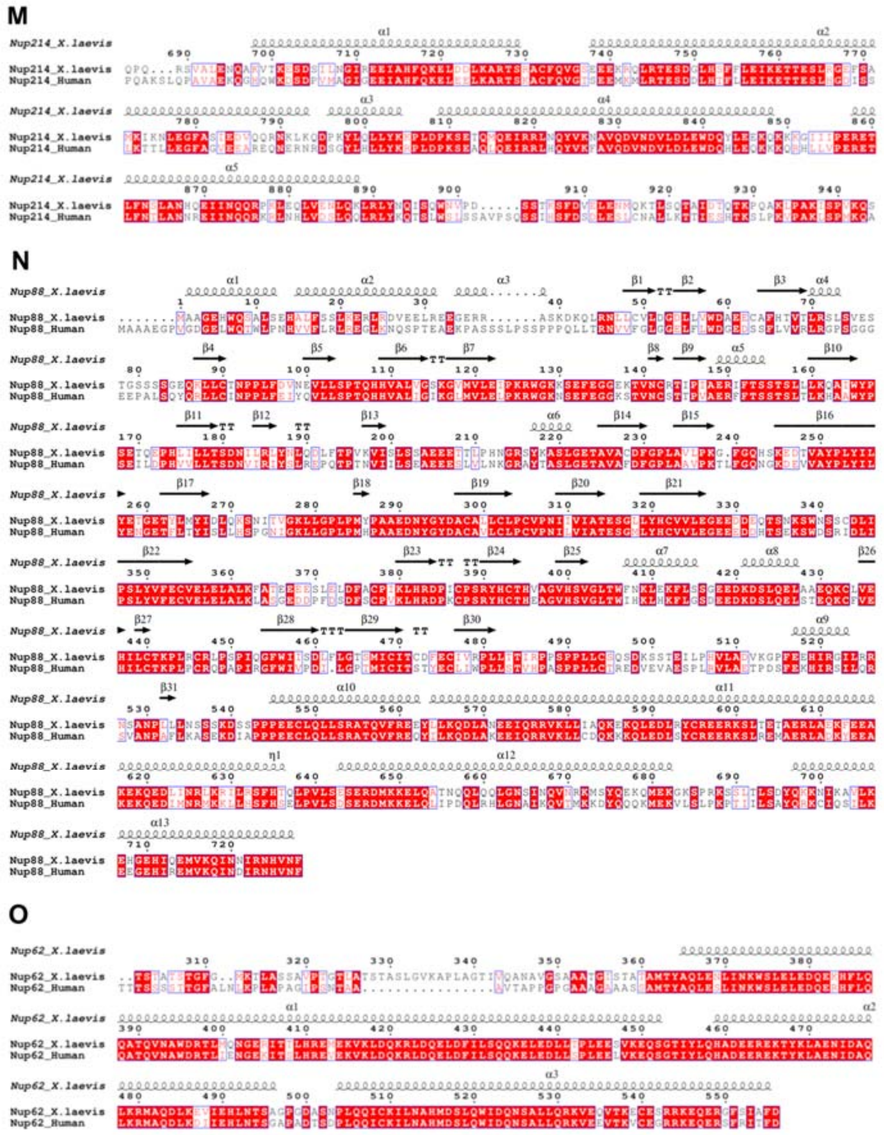


**Extended Data Figure 8. Sequence alignment of CR Nups from *X. laevis* and human NPCs. (A-O)** Nup85, Nup43, Seh1, Nup160, Nup37, Nup96, Sec13, Nup107, Nup133, Nup205, Nup93, Nup358, Nup214, Nup88 and Nup62, respectively. Secondary structures are also indicated.

**Extended Data Table 1. Statistics of cryo-SPA data collection and image processing.**

| | | | |
|---|---|---|---|
| **Data acquisition** | | | |
| Microscope | Titan Krios G2 | | |
| Voltage (kV) | 300 | | |
| Detector | Gatan K2 | | |
| Energy filter | Gatan GIF Quantum, 20 eV | | |
| Mode | Super resolution | | |
| Pixel size (Å) | 2.24 | | |
| Stage tilting angle | 30°/ 45°/ 60°/ 0°(side-view) | | |
| Exposure per tilt (e/Å²) | 60 / 80/ 100/ 100 (or 120) | | |
| Number of images | 8745 | | |
| Defocus range (μm) | -1 ~ -4 | | |
| Software | SerialEM | | |
| **Reconstruction** | | | |
| Software | RELION-3.0/CryoSPARC | | |
| Data set | CR core region | CR subunit region | CR Nup358 region |
| Final number of particles | 354460 | 354460 | 678866 |
| Symmetry | C1 | C1 | C1 |
| Final resolution (Å) | 8 | 8.7 | 8.9 |
| Map pixel size (Å) | 2.24 | 2.24 | 2.24 |
| Map sharpening B-factor (Å²) | -591 | -763 | -850 |

### Supplementary information

**Supplementary Video 1.**

Model-map fitting quality of CR Nups from the *X. laevis* NPC.
